# Learning torus PCA based classification for multiscale RNA backbone structure correction with application to SARS-CoV-2

**DOI:** 10.1101/2021.08.06.455406

**Authors:** Henrik Wiechers, Benjamin Eltzner, Kanti V. Mardia, Stephan F. Huckemann

## Abstract

**Motivation:** Reconstructions of structure of biomolecules, for instance via X-ray crystallography or cryo-EM frequently contain clashes of atomic centers. Correction methods are usually based on simulations approximating biophysical chemistry, making them computationally expensive and often not correcting all clashes.

**Results:** We propose a computationally fast data-driven statistical method yielding suites free from within-suite clashes: From such a clash free training data set, devising mode hunting after torus PCA on adaptive cutting average linkage tree clustering (MINTAGE), we learn RNA suite shapes. With classification based on multiscale structure enhancement (CLEAN), for a given clash suite we determine its neighborhood on a mesoscopic scale involving several suites. As corrected suite we propose the Fréchet mean on a torus of the largest classes in this neighborhood. We validate CLEAN MINTAGE on a benchmark data set, compare it to a state of the art correction method and apply it, as proof of concept, to two exemplary suites adjacent to helical pieces of the frameshift stimulation element of SARS-CoV-2 which are difficult to reconstruct. In contrast to a recent reconstruction proposing several different structure models, CLEAN MINTAGE unanimously proposes structure corrections within the same clash free class for all suites.

**Code Availability:** https://gitlab.gwdg.de/henrik.wiechers1/clean-mintage-code

## 1 Introduction

Understanding the structure of active biomolecules is ever more important for maintaining and prolonging human health (e.g Schlick and Pyle (2017)). In particular this pertains to RNA molecules in view of designing drugs that target specific structures (e.g. Batool *et al*. (2019)), as recently impressively demonstrated by the worldwide effort confronting the SARS-CoV-2 (severe acute respiratory syndrome) virus responsible for the COVID-19 (corona virus disease) pandemic (e.g. Croll *et al*. (2021)).

Extracting RNA primary structure is nowadays fairly well feasible using currently available gene sequencing technology (e.g. Wang *et al*. (2009)). Predicting from that the structure, however, is a still unsolved fundamental problem (e.g. Schlick and Pyle (2017)). Although elaborate methods such as X-ray crystallography and cryo-EM (cryogenic electron microscopy) are used that return spatial electron densities – and from these densities individual atom positions can be inferred – frequently, the inferred molecular structures contain so-called clashes (e.g Murray *et al*. (2003); Schlick and Pyle (2017)).

#### Definition 1.1.

*A* **clash** *is a forbidden molecular configuration, where two atoms are reconstructed closer to each other than is chemically possible*^1^.

In case of RNA, clashes most relevant and most difficult to correct are between atoms along the backbone, in particular when single hydrogen atoms are added to inferred structures (e.g. Murray *et al*. (2003)).

In order to correct such clashes, methods from *molecular dynamics* are usually employed: Simulated atoms are allowed to fluctuate into positions of minimal energy, following approximations of the laws of biophysical chemistry (e.g. Chou *et al*. (2013)). For RNA molecules, these simulations are highly computation intensive due to the large variability of RNA shape. If local and not global energy minima are achieved, thus corrected molecules may still feature clashes and their geometries may be outliers in comparison to clash free geometries (e.g. Richardson *et al*. (2018)). Figure 2 (center) illustrates the state of the art correction by ERRASER (Enumerative Real-space Refinement ASsisted by Electron-density under Rosetta) from Chou *et al*. (2013) and (right) our CLEAN MINTAGE (Classification based on muLtiscale structurE enhAncemeNt using Mode huntINg after Tours pca on Adaptive cutting averaGe linkage trEes) correction proposed here. As our method is developed from a statistical learning perspective it does not suffer from the above drawbacks as evident from Figure 2.

**Fig. 1:**
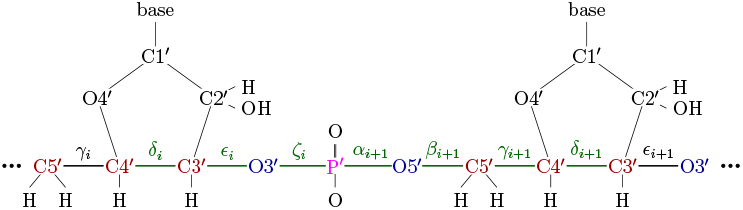
Backbone suite number i with 7 dihedral angles δ_i_,ε_i_, ζ_i_, α_i +1_, β_i+1_, γ_i+1_, δ_i+i_ describing the suite’s shape, see Figure 17 in the supplement.

**Fig. 2:**
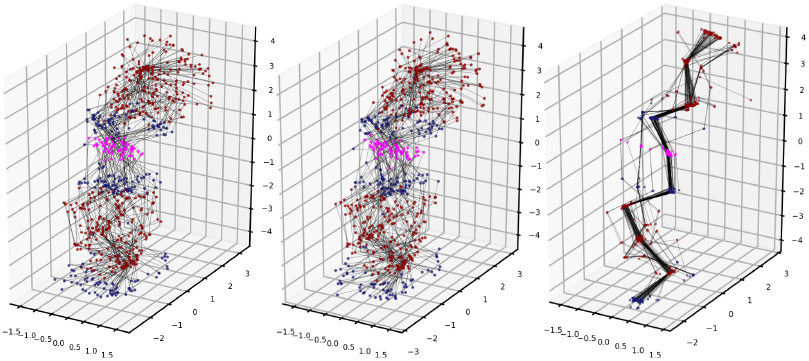
RNA backbone suites with carbon (dark red), oxygen (dark blue) and phosphorus atoms (pink), see Figure 1. Left: 73 clash suites which form our ERRASER test set 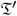 (see Supplement I). Center: Their corrections by ERRASER. More than half of them still feature clashes (see Supplement I.1). Right: Their corrections by CLEAN MINTAGE. Each correction belongs to a clash free class. See Figures 14 and 15 in the supplement for both corrections in dihedral angle representation.

Our method is computationally fast and proposes data-driven corrections taken from classes of clash free geometries previously learned using *torus PCA* from Eltzner *et al*. (2018). It draws power, on the one hand from a specifically designed adaptive pre-clustering method and the ability of torus PCA to efficiently reduce torus clusters to one dimension only, which then allows for further post clustering by mode hunting (Figure 3) and, on the other hand from the simultaneous two-scale approach, inspired by Hamelryck *et al*. (2010) using “coarse and fine degrees of freedom”. It may well serve to provide clash free initial configurations for subsequent molecular dynamics methods.

**Fig. 3:**
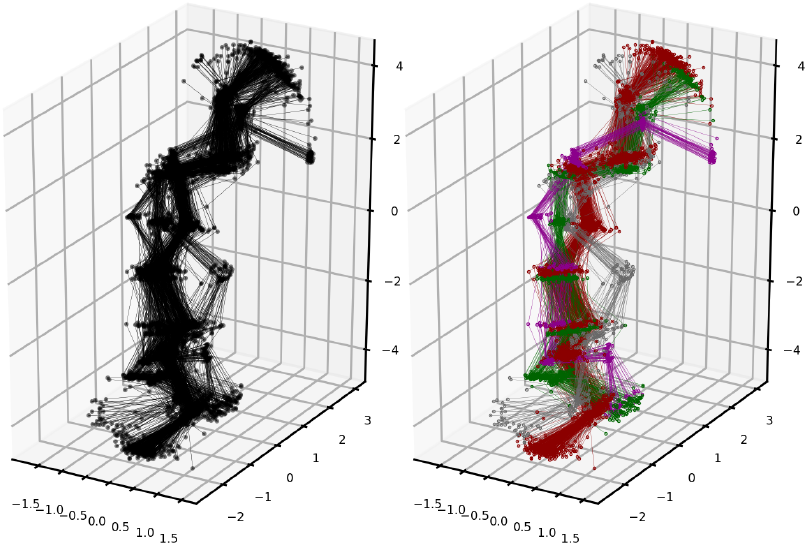
One-dimensional torus PCA representation of torus pre-cluster 2 from the benchmark training set (Section G.2) ofRNA suite backbone shapes (left) is efficiently decomposed into 4 classes (right) by mode hunting, yielding Class 2 (red), Class 5 (green), Class 6 (grey), Class 7 (purple), see also Figure 20 in the supplement.

As most RNA backbone clashes appear within ***suites*** (the part from one sugar ring to the next (e.g. Murray *et al*. (2003), see Figure 1 and Definition D.1 in Supplement D), in our ***microscopic*** scale we represent suites by tuples of 7 dihedral angles. Recall that every directed bond between two atoms (C4’ and C3’, say, cf. Figure 1 and Figure 17 in the supplement) defines a *dihedral angle* in the one-dimensional *torus* 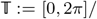 (here “~” identifies 0 and 2π) with distance 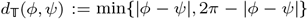, 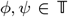. Hence, a suite’s shape is a data point on the seven dimensional torus 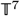 with the canonical product distance 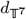 given by

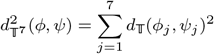

with 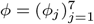, 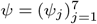

#### Definition 1.2.

*We call a suite a* **clash suite** *if two of its backbone atoms (including associated hydrogen atoms and oxygen atoms belonging to the phosphate) clash with each other. All other suites that have k (specified below) neighboring non-clash suites are called* **clash free**.

Using our Algorithm 2.1 (MINTAGE), we learn 15 classes of clash free suites including their variability from a large clash free benchmark training data set, see Figure 3, called the ***MINTAGE benchmark classes***.

Our ***mesoscopic*** scale comprises for every suite, whether clash free or a clash suite, *k* neighboring suites on either side. This gives a configuration of 2*k* + 1 suites, representing the local geometry of the suite’s neighborhood. Setting *k* = 2 the mesoscopic scale of a suite within a helix will conveniently correspond roughly to a half turn. We describe the *mesoscopic shape* by the *size-and-shape* of the centers of the 6 = 2*k* + 2 sugar rings involved (Figure 4), giving an element in 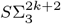 with the *Procrustes distance d_Σ_*, see Supplements C, D.3 and Dryden and Mardia (2016).

**Fig. 4:**
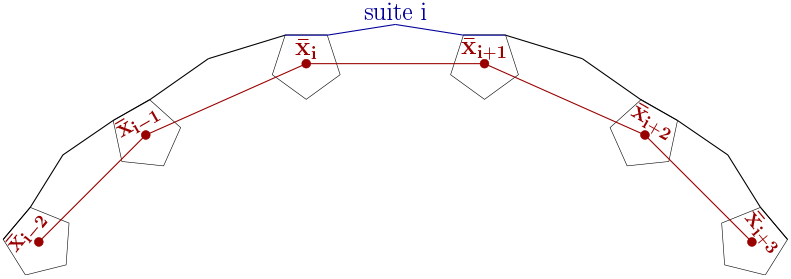
The mesoscopic shape (red lines) for k = 2 centered at the i-th suite is determined by the six centers of the sugar rings 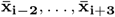. Their connecting backbones (blue and black lines) give 5 suites, two before and two after suite i.

Our Algorithm 2.4 (CLEAN) corrects a clash suite as follows. With respect to the geometry of mesoscopic shape space, its *ρ* (we use *ρ* = 50) nearest clash free neighbors are determined and the class containing most of the corresponding *ρ* suites is identified, defining a subset of the *ρ* suites. Then, the clash suite is replaced by the torus Fréchet mean of the subset of suites, yielding a clash free suite in a data-driven manner. The mesoscopic shape of the clash suite is replaced with the orthogonal projection of the Procrustes mean of the subset’s mesoscopic shapes to the subset of mesoscopic shapes featuring specific constraints inherited from the mesoscopic shape of the clash suite.

When we apply our method to two exemplary suites adjacent to helical structures of the *frameshift stimulation element* of SARS-CoV-2 (detailed in Section 3.2), our method proposes only elements from a single class against various different structure proposals in Zhang *et al*. (2020), see Figure 5.

**Fig. 5:**
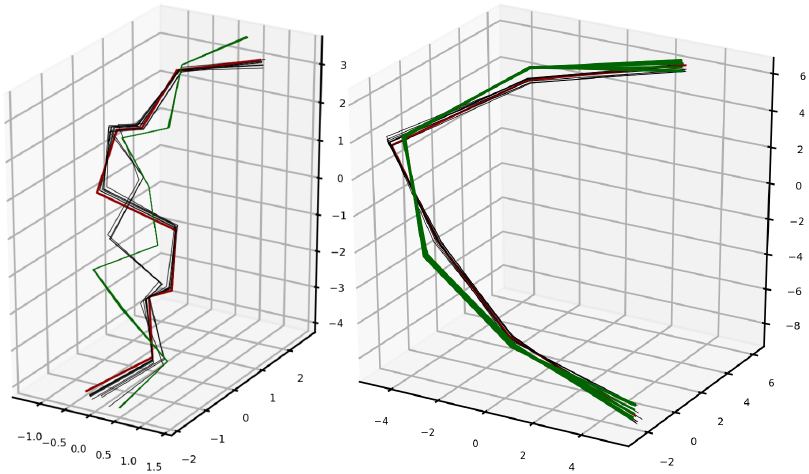
Left: Ten different model proposals by Zhang *et al*. (2020) for the backbone of Suite 2 (from the left panel of Figure 11 in Section 3.2); adjacent to a helical piece of the frameshift stimulation element of SARS-CoV-2. One model (red) clashes (with clash score 0.401Å which is slightly above the threshold of 0.4Å) and nine models (black, featuring two different types of shape) are clash free, but inconsistent. Our CLEAN MINTAGE correction proposes unanimously over all ten models a suite shape (green, all consistent, so they overlap) from the clash free Class 1 (see Figures 13 and 20 from the supplement). Right: Mesoscopic shapes of the ten models (red and black) and their ten CLEAN MINTAGE corrections (green).

For graphical illustration we display torus-valued shapes also as size-and-shape configurations. In displays, all size-and-shape configurations will be shown in optimal position to the Procrustes mean of the underlying data set.

Size-and-shape space and Fréchet means are detailed in the supplement, along with details on our two scales, reconstruction methods, as well as further details on our methods and results. In particular the benchmark data sets are detailed in Supplement D.4.

## 2 Methods

### 2.1 Torus PCA Based Clustering

Our *Mode huntINg after Torus pca on Adaptive cutting averaGe linkage trEes* (MINTAGE) algorithm builds on three components. Initially the torus data is pre-clustered by adaptively cutting an average linkage clustering tree, as detailed in Supplement G.1. Then, each cluster is reduced to a one dimensional torus representation using *torus PCA*, recently developed by Eltzner *et al*. (2018), with varying flag parameters. While the torus is rather inconvenient for PCA based dimension reduction methods (almost all geodesics are dense and tangent space methods lose periodicity), torus PCA deforms a torus into a stratified sphere, opening up the toolbox of *principal nested spheres* from Jung *et al*. (2012). This makes the torus even more attractive for PCA based dimension reduction methods than Euclidean space, cf. Huckemann and Eltzner (2015). Finally, if the sum of squared residual distances to these one dimensional torus representations of the pre-clusters is less than one fourth of the cluster’s total Fréchet variance, these one dimensional representations are subjected to circular mode hunting that identifies subclusters with statistical significance, as detailed in Section 2.2.

#### Algorithm 2.1 (MINTAGE)

1. From input suite data *X*^(1)^,…, *X*^(*n*)^ obtain a list of pre-clusters using Algorithm G.1 from the supplement and store it as the *remaining cluster list* 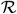. Create the initially empty *final cluster list* 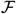.
2. While 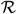 is non-empty:

a. Take a cluster *C* from 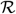 and set *m* = 1:
b. For *C* perform torus PCA from Eltzner *et al*. (2018) with the flags GC (gap centered), MC (mean centered), SI (spread inside), SO (spread outside) as detailed there

- if m = 1: GC, SI
- if m = 2: GC, SO
- if m = 3:MC, SI
- if m = 4:MC, SO
- if m = 5: Remove *C* from the remaining cluster list 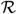, add it to 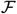 and go to Step 2(a).
c. For the suite *X*^(*j*)^ ∈ *C* let *X* ^(*j*,1*D*)^ denote their one-dimensional torus PCA projections and let *μ* denote the torus PCA nested mean from Eltzner *et al*. (2018). Whenever 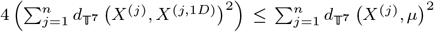, perform mode hunting from Section 2.2:

- if subclusters were found, add them to the remaining cluster list 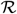 and remove *C* from it.
- else: Set *m* = *m* + 1 and go to Step 2(b).
3. Return 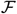.

Supplement G.2 details the application of MINTAGE to the benchmark training data set yielding ***the 15 MINTAGE benchmark classes*** and ***the MINTAGE benchmark outliers***.

### 2.2 Circular Mode Hunting

We cluster the one-dimensional projections obtained by torus PCA in Step 2(c) of Algorithm 2.1 using the multiscale method described by Dümbgen and Walther (2008). Although this method was originally defined for the real line, its numerical implementation for circular data is even simpler. Since modes are separated by minima, we use this method to identify regions in which minima are located with a certain confidence level. (Throughout the applications, we use a fixed confidence level of 95%). In each of these regions, we estimate the minima by estimating the density of the one-dimensional projections using a wrapped Gaussian kernel. As the bandwidth increases, the number of minima of the density estimate in each of the regions decreases. Whenever there is only one minimum left in a region – this will inevitably happen due to the causality of the wrapped normal kernel, see Huckemann *et al*. (2016) – we take this as a cluster boundary, see Figure 6 for an illustration of our method. Figure 3 shows pre-cluster 2 from the benchmark training data set which is decomposed by mode hunting into four subclusters.

**Fig. 6:**
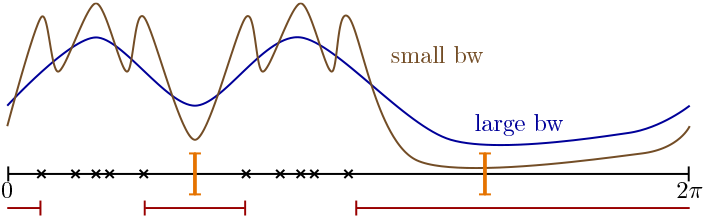
Circular mode hunting for data (black asterisks). Intervals containing minima with statistical significance (red) and wrapped Gaussian kernel smoothed densities with varying bandwidths having too many minima (brown) and having the statistically significant number of minima (blue). The latter minima are taken as cluster boundaries (orange).

### 2.3 Multiscale RNA backbone correction

We first motivate why and how our multiscale correction algorithm involves the two different scales of suites and mesoscopic shapes. Moreover, in Supplement H.2 we validate it on the benchmark test data set from Supplement D.4.

#### 2.3.1 Motivation for a Multiscale Correction Ansatz

In a first fundamental study, we establish a relationship between suites from the benchmark training data set (Supplement D.4) that have similar mesoscopic shapes, see Figure 4 and Definition D.2 in Supplement D.3. To this end, we cluster the mesoscopic shapes of the suites from the benchmark training data set using the *simple version* of Algorithm G.1 from Supplement G.1, performing only Steps 1 and 2 with *κ* = 5 and *d*_max_ such that 50% of the mesoscopic strands are outliers. This guarantees that the *simple mesoscopic clusters* obtained are rather concentrated and for this reason we obtain a higher (than in Supplement G.2) number of 110 such clusters. It turns out that the suites corresponding to each simple mesoscopic cluster also form rather concentrated suite clusters and they are in high correspondence with the 15 MINTAGE benchmark classes (obtained from the MINTAGE Algorithm 2.1 applied in Supplement G.2 to the benchmark training set). This leads to the following hypothesis.

#### Hypothesis 2.2.

*Correctly reconstructed suites with similar mesoscopic shapes have also similar suite shape. In particular, concentrated mesoscopic clusters relate to suite clusters*.

Indeed, for the overwhelming majority of the simple mesoscopic clusters the standard deviation of angles (between 0 and 2π) of their suites is less than 0.6, as depicted in the left panel of Figure 21 in the supplement. Only very few clusters with low cardinality (close to the minimum of *κ* + 1 = 6) have higher suite variance. On the other hand, the high correspondence of simple mesoscopic clusters with the MINTAGE benchmark classes is detailed in the caption of Figure 7 and in the right panel of Figure 21 in Supplement H.1.

**Fig. 7:**
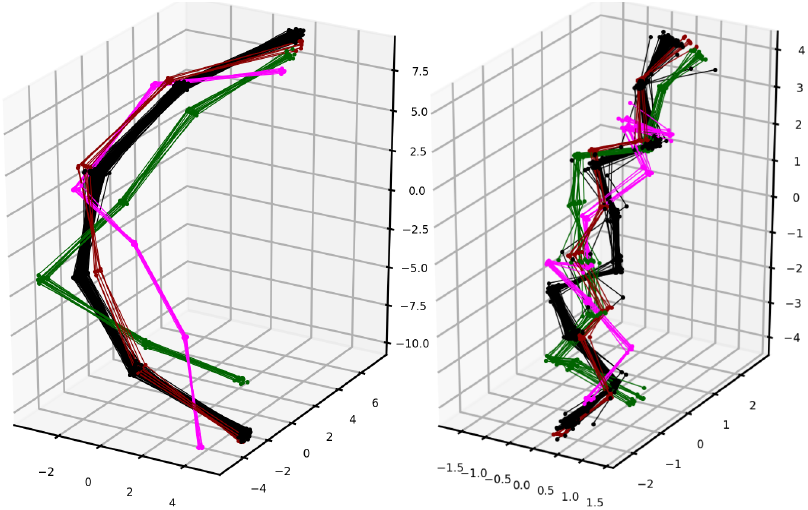
Left: Exemplary *simple* mesoscopic clusters. Right: Suites belonging to the mesoscopic shapes from the exemplary clusters. Notably, Cluster 1 (black) of size 77 contains 73 suites from MINTAGE benchmark Class 1, all of the others clusters are in 1-to-1 correspondence to MINTAGE benchmark classes: Cluster 30 (green, size 13) to Class 5, 55 (magenta, size 8) to 12 and 92 (red, size 6) to 2.

In a second fundamental study, we consider the 198 clash suites in the benchmark data set forming the test data set, see Supplement D.4. Their suite shapes as well as their mesoscopic shapes feature a rather larger spread, see Figure 22 in the supplement. As before, we consider training suites from concentrated neighborhoods in the mesoscopic shape space, of size 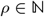. For a given clash suite 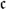 such a neighborhood is

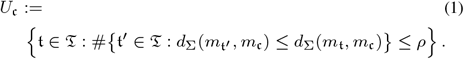

Here 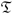 is the set of training suites (the clash free suites in the benchmark data set, see Supplement D.4) and mt denotes the mesoscopic shape of 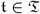. From Figure 8, which illustrates a typical clash suite 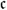, we derive the following hypothesis.

**Fig. 8:**
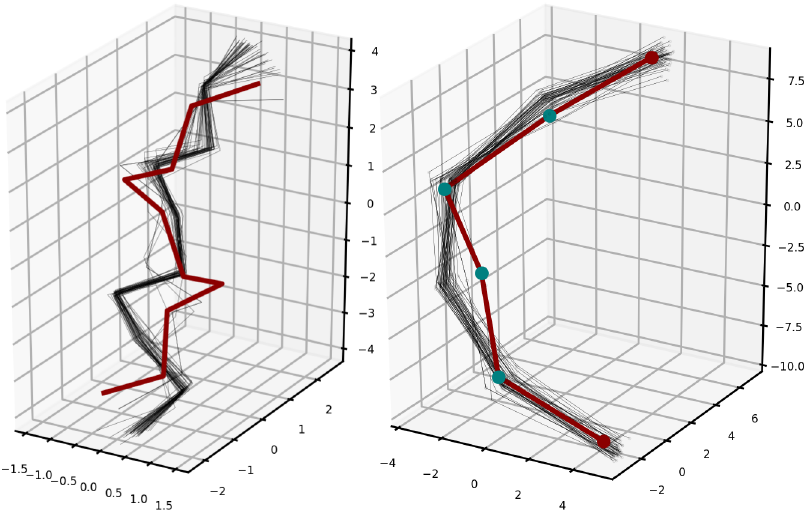
Left: A typical clash suite 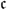 (red) and the *ρ* = 50 suites (black) from the neighborhood 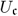 in the benchmark training data set with respect to mesoscopic shape distance. Right: The corresponding mesoscopic shape 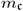 (red) and those from 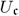 (black). The latter are concentrated in the mesoscopic shape space.

##### Hypothesis 2.3.

*While clash suite shapes are rather irregular among the suite shapes of their clash free neighbors in mesoscopic space (left panel), their mesoscopic shapes differs mostly mildly from nearby clash free mesoscopic shapes (right panel)*.

In particular, two of the four teal landmarks in the middle differ systematically from those of the clash free mesoscopic shapes, thus hinting also towards wrongly reconstructed sugar ring centers. Again, 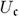 is rather concentrated in the mesoscopic shape space and its suites feature a strongly dominating cluster. For this reason, the following multiscale backbone correction algorithm simultaneously corrects clashing suites on suite and on mesoscopic levels, working with concentrated neighborhoods defined by mesoscopic shape distance. In these concentrated neighborhoods, dominating clusters from MINTAGE of the training data set provide guidance for correction.

#### 2.3.2 The RNA Backbone Correction Algorithm

We exploit the above Hypotheses 2.2 and 2.3, take into account that the suite of interest in the mesoscopic shape is positioned between the third and fourth sugar ring, and hence a small change in one of the center sugar rings of the mesoscopic shape can have a greater impact on the shape of the suite in the center. The design of the algorithm is further discussed in Remark H.2 from Supplement H.1.

#### Algorithm 2.4 (CLEAN). Input

- a training data set 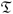 comprising only non clashing and admissible suites, i.e. suites that come along with a mesoscopic shape, see Definition D.4 in the supplement,
- a cluster list *C*_1_,…, *C_r_* and an outlier set *R* for 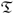 obtained from applying the MINTAGE Algorithm 2.1,
- a tuning parameter 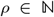 governing the size of neighborhoods defined via mesoscopic shape distance, we choose *ρ* = 50, and
- a clash suite 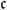 and its corresponding mesoscopic clash shape 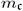.
- the flag dominating set to absolute or relative which will return either the absolutely dominating cluster 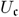 from (1) or the relatively dominating cluster with at least *ρ*/10 elements, taking into account cluster size.

Algorithm steps:

1. Calculate

a. the neighborhood 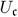 as defined in (1) of the *ρ* suites of the training data, whose mesoscopic shapes are most similar to 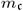 with respect to mesoscopic shape space distance;
b. according to flag dominating, the number 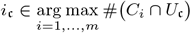, (absolute), or, 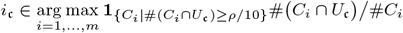, (relative), respectively, of the dominant torus PCA cluster in 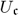;
c. a Fréchetmean 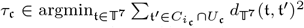, of the dominant cluster’s suites in the neighborhood;
d. the approximate length 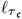 of the suite by the mean distance of the two C4 atoms in the suites of 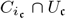;
e. a Procrustes mean

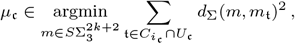

of the corresponding mesoscopic shapes,
2. With a mesoscopic shape 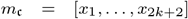 defined as in Equation (2) in the Supplement by a representative (*x*_1_, …, *x*_2*k*+2_), determine with the Procrustes type Subalgorithm H.1 from Supplement H.1 the corrected mesoscopic shape 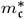 as the orthogonal projection *Y*′ of the Procrustes mean 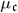 to the set

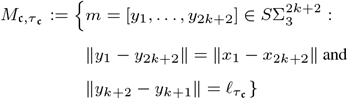

of mesoscopic shapes whose configurations share the first and the last landmark with a configuration of 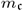 and whose distance between the central landmarks is the length 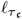 of the configuration Fréchet mean suite 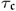.
3. Return: the corrected suite shape 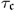 and its corrected mesoscopic shape 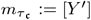.

For all applications we used DOMINATING = ABSOLUTE. Figure 9 and Figure 15 in the supplement display also DOMINATING = RELATIVE.

**Fig. 9:**
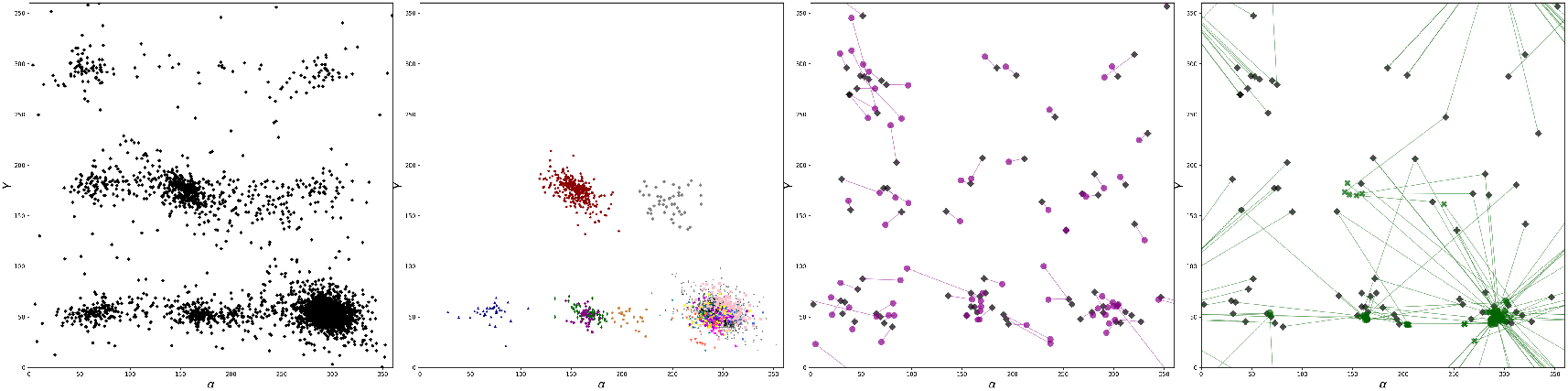
Dihedral angles α (horizontal) and γ (vertical). Left: All suites from the benchmark training data set. Centre left: All MINTAGE classes (without outliers). Centre Right: 73 clash suites from the benchmark test data set (black) and their ERRASER corrections (magenta). Right: Same clash suites (black) and their CLEAN MINTAGE corrections by Fréchet means of the dominating class (green circles for DOMINATING = ABSOLUTE and green crosses for DOMINATING = RELATIVE). Figures 12, 13, 14 and 15 in the supplement show the situation for all dihedral angle combinations.

## 3 Results

### 3.1 Comparing CLEAN MINTAGE with ERRASER

Applying molecular dynamics with the clash suite as initial state, ERRASER corrects clash suites locally and moderately. This often results in proposals that remain clashing and outlying. In contrast, corrections by CLEAN MINTAGE are guided by minor changes on the mesoscopic scale, which may yield more drastic suite corrections. These are, however, clash free and no longer outlying, as illustrated in Figures 9 and 10, and detailed in Supplement I.

**Fig. 10:**
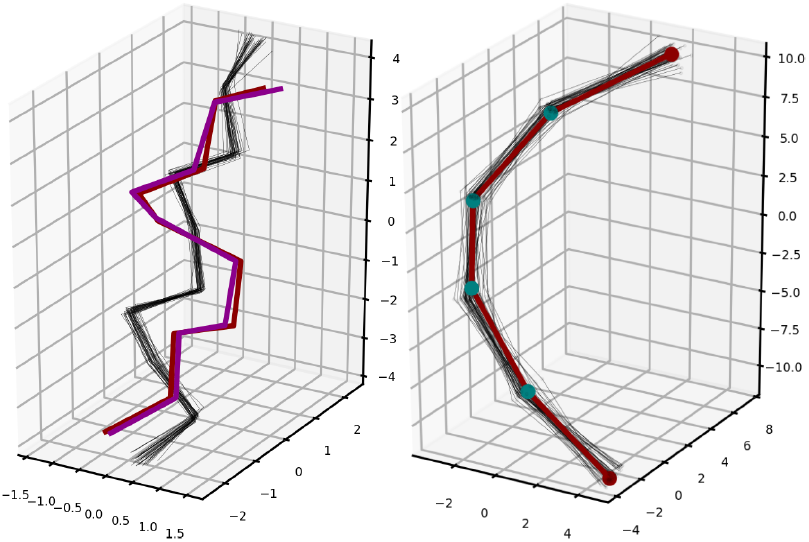
Left: A clash suite (red) is only infinitesimally corrected by ERRASER (magenta), remaining clashing and outlying. The more drastic CLEAN MINTAGE correction (the Fréchet mean of the black suites constituting the main cluster of the neighborhood 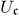 from (1)) is guided by a minimal correction of the mesoscopic shape of the clash suite. Right: The mesoscopic shape of the clash suite (red) lies almost in the middle of the mesocopic shapes corresponding to the MINTAGE cluster, heavily dominating its local neighborhood 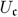 (black, 39 out of 50 suites).

### 3.2 COVID 19 Backbone Correction

With the recent worldwide pandemic of the *severe acute respiratory syndrome coronavirus 2* (SARS-CoV-2), the virus’ RNA structure reconstruction and backbone correction has become ever more relevant. Indeed, effective drug and vaccine development necessitates good understanding of the three-dimensional RNA structure, see Croll *et al*. (2021). Recently, a large number of measurements has been added to the Protein Data Bank, see Berman *et al*. (2000), and as part of the *Coronavirus Structural Task Force* (CSTF) headed by Andrea Thorn, a large number of data sets of SARS-CoV-2 and related structures are compiled in a git repository, see Thorn *et al*. (2021). While X-ray crystallography can achieve very high resolution in principle, the large viral genome, comprising ~ 20, 000 bases, is very difficult to crystallize. Therefore, many structures are determined by cryogenic electron microscopy (cryo-EM).

#### 3.2.1 The Frameshift Element

In Zhang *et al*. (2020), the *frameshift stimulation element* (FSE) of the SARS-CoV-2 genome was studied, which, due to its *slippery site* encodes different proteins simultaneously. As their balanced expression is required for virus replication, this element is believed to be fairly resistant against mutations. Hence it is a promising target for antiviral drug design. Its three-dimensional structure has been assessed by cryo-EM with a resolution of 6.9Å using the ribosolve pipeline from Kappel *et al*. (2020). Using a consensus secondary structure of the molecule and the cryo-EM map, Zhang *et al*. (2020) proposed 10 possible threedimensional structure models (based on a measurement with mean pairwise root mean squared deviation of 5.689Å) and stored them to the Protein Data Bank. Notably, it was not possible to reliably assign individual atom positions, but the secondary arrangement of helical segments and the non-helical linking segments could be reconstructed,see Zhang *et al*. (2020) and first panel of Figure 11. In particular, the suites linking different helical segments have been difficult to reconstruct. Here we focus on Residues 28/29 which we call *Suite 1* and on Residues 33/34 which we call *Suite 2* (referring to enumeration in the PDB file).

**Fig. 11:**
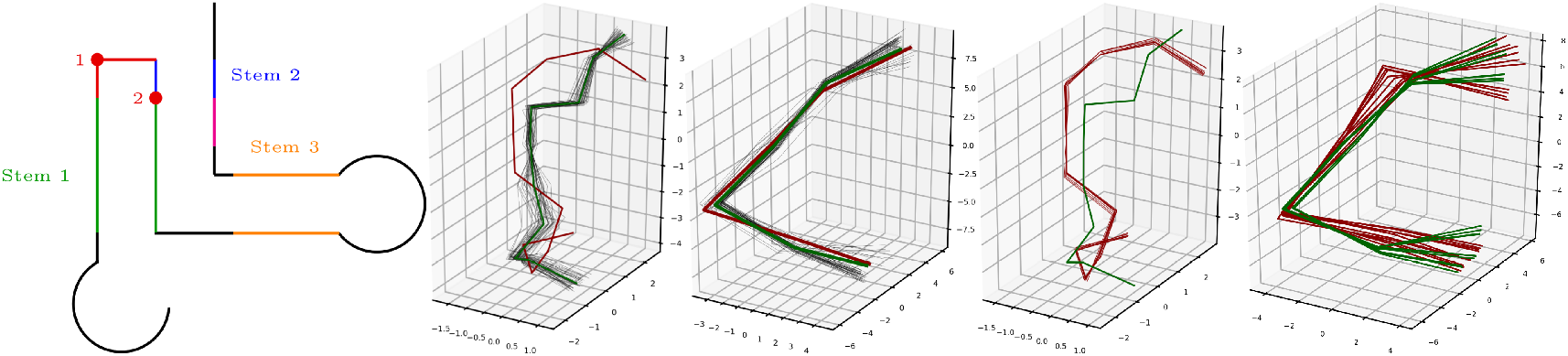
First: Secondary structure of the imputed frameshift stimulation element, cf. (Zhang *et al*., 2020, Figure 8). The green, blue and orange regions show the double-stranded helix regions. The red dots (only one nucleotide is on the red branch) between depict clashing Suites 1 and 2. Second: Model 1 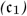 of Suite 1 (red, clashing) proposed by Zhang *et al*. (2020), the 43 suites from Class 5 in neighborhood 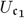 (black) and their torus mean (green). Third: The corresponding mesoscopic shape 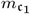 (red), the 43 mesoscopic shapes of suites from Class 5 in neighborhood 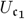 (black) and their Procrustes mean (green). Fourth: All models 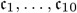 (red, clashing) proposed by Zhang *et al*. (2020) and their CLEAN MINTAGE corrections (green). Fifth: Mesoscopic shapes 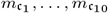 (red) of the 10 models and projections of Procrustes means (green) of respective dominating clusters.

Suite 1 (one of the two red dots in the left panel of Figure 11) is a clash suite in all 10 models, as determined by PHENIX, see Supplement E. Moreover, the P’-O3’ bond is unphysically long (red vertical in 2nd and 4th panel of Figure 11), reinforcing the conclusion of a bad structure fit. With the MINTAGE benchmark classes from Supplement G.2, CLEAN yields 10 corrections, one for each of the 10 models.

The fourth panel of Figure 11 shows the clash suites 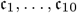, from each of the models. These suites are very similar and so are their mesoscopic shapes in the fifth panel. In every neighborhood 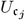 (*j* = 1,…, 10) constructed by CLEAN, MINTAGE benchmark Class 5 from Supplement G.2, see Figure 20 in the supplement, is strongly dominating. The means of the suites of Class 5 in the respective neighborhoods are nearly identical and give the universal correction for Suite 1, also displayed in the fourth panel of Figure 11. The second panel of Figure 11 shows the closeup for clash suite 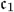. In 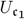, 43 out of the 50 suites belong to Class 5.

Suite 2 (at the other red dot in the first panel of Figure 11) is a clash suite only for Model 5, the suites of all other 9 models are clash free, but inconsistent (see Figure 5). Moreover, in the other models for Suite 2 there are other clashes, such as clashes between backbone and base atoms. Again with the MINTAGE benchmark classes from Supplement G.2, CLEAN yields 10 corrections, one for each of the suites 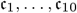 from the 10 models. As before, the corresponding mesoscopic shapes are very similar, see right panel of Figure 5, and there is a single MINTAGE benchmark class that dominates every neighborhood, namely Class 1. In consequence, the CLEAN MINTAGE correction is the torus mean of the part of Class 1 in the respective neighborhoods. Again these are very similar, giving the universal CLEAN MINTAGE correction for Suite 2, now depicted (green) in the left panel of Figure 5.

## 4 Discussion

The RNA backbone structure correction method proposed in this paper rests on two pillars:

1. Torus PCA based suite classification of a large benchmark clash free data set and
2. multiscale suite correction based on proximity on a mesoscopic scale.

(1): Remarkably, approximately one out of 6 (see Figure 20 in the supplement) of the benchmark data suites (and approximately one out of 7 of the ERRASER training suites, see Figure 26 in the supplement) could not be classified by MINTAGE. While there may be multiple reasons for this, one is rather obvious: As the presence of clashes is a mere indicator for faulty atom configuration proposals, it is very likely that many clash free outliers are also due to faulty reconstructions and hence deserve structure correction as well. Since our method is specifically designed to correct suites in such a way that their shapes resemble abundant classes of shapes, it is well suited to reduce outliers also in cases where no clash occurs.

(2): Our approach at multiple scales, at a specific suite and at a specific mesoscopic scale (of roughly one half helix turn length) has proved to be successful for correcting clash suites via assignment to clash free classes. A more complete picture of the improved structure may be achieved by simultaneously correcting several suites along the following lines:

1. A central suite correction may warrant simultaneous correction of the suites involved at the mesoscopic scale (notably, adjacent suites overlap at four atoms),
2. Whenever there are base bindings from a corrected suite to other suites, their structures may be simultaneously corrected.
3. Having thus far corrected only within-suite backbone clashes, which are very common, a further refinement aims at simultaneously correcting different suites that clash with one another.
4. Notably, the multiscale correction method can be applied, potentially, to various other biomolecules and in particular to protein structure correction, see Hamelryck *et al*. (2010).

At this point we would like to clarify that ERRASER corrects not only all suites (clashing and clash free), it also takes into account potential clashes with atoms from more distant suites and their nucleobases and performs various other structure improvements. While this entire process took several days on the ROSIE servers Chou *et al*. (2013) our CLEAN MINTAGE, only correcting within suite clashes, ran within minutes. Recall that ERRASER, due to its moderate correction potential based on molecular dynamics, was only able to completely remove clashes for less than half of the benchmark data. While our outputs, yielding corrections within clash free classes, do not provide a comprehensive treatment, for instance, leaving neighboring suites unchanged, they may serve as initial states for subsequent molecular dynamics, and thus provide powerful guidance.

## Acknowledgements

The authors are grateful for fruitful discussions with Markus Hiller, Thomas Hamelryck and Carina Wollnik.

## Supplementary Information

### A PDB files

Table 1 lists the PDB IDs and the corresponding resolution of the benchmark data set, see Supplement D.4.

**Table 1.**
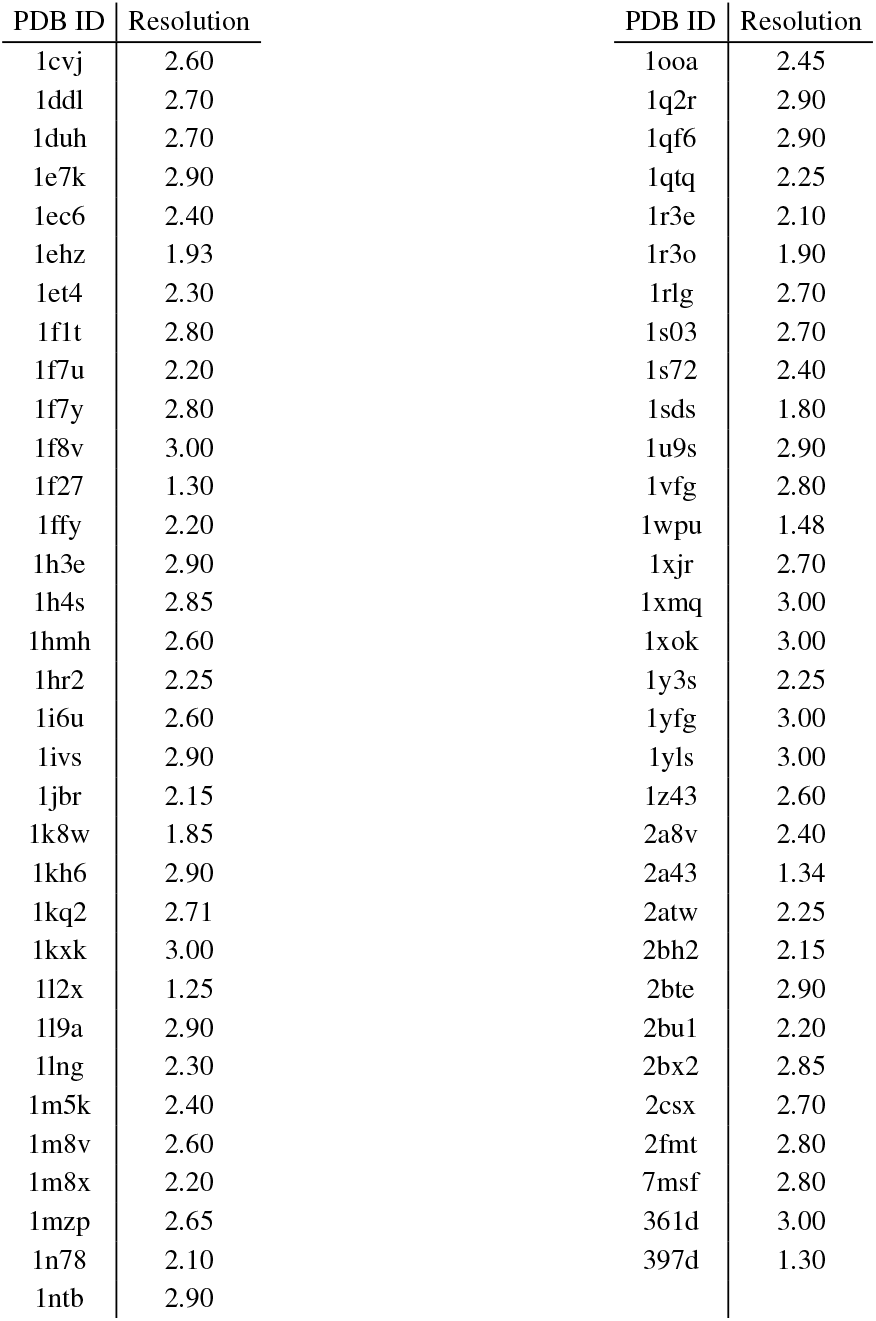
The PDB IDs from the 71 different measurements from the benchmark data set, see Supplement D.4.

### B Scatterplots of all pairwise dihedral angle combinations

Figure 12 illustrates all two dimensional dihedral angle combinations of the suites of the training data set 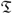, introduced in Supplement D.4. Their MINTAGE classes (see Algorithm 2.1 in the main text) with the parameters described in Supplement G.2 are shown in Figure 13. The corresponding cluster sizes are summarized in Figure 20. The scatterplots of the 73 clash suites 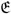 from the ERRASER test set are depicted in Figures 14 and 15 as black diamonds. Figure 14 shows the corresponding ERRASER corrections as magenta circles (see Supplement I) and Figure 15 shows the corresponding CLEAN MINTAGE (Algorithm 2.4 in the main text) correction as green circles and crosses, respectively.

**Fig. 12:**
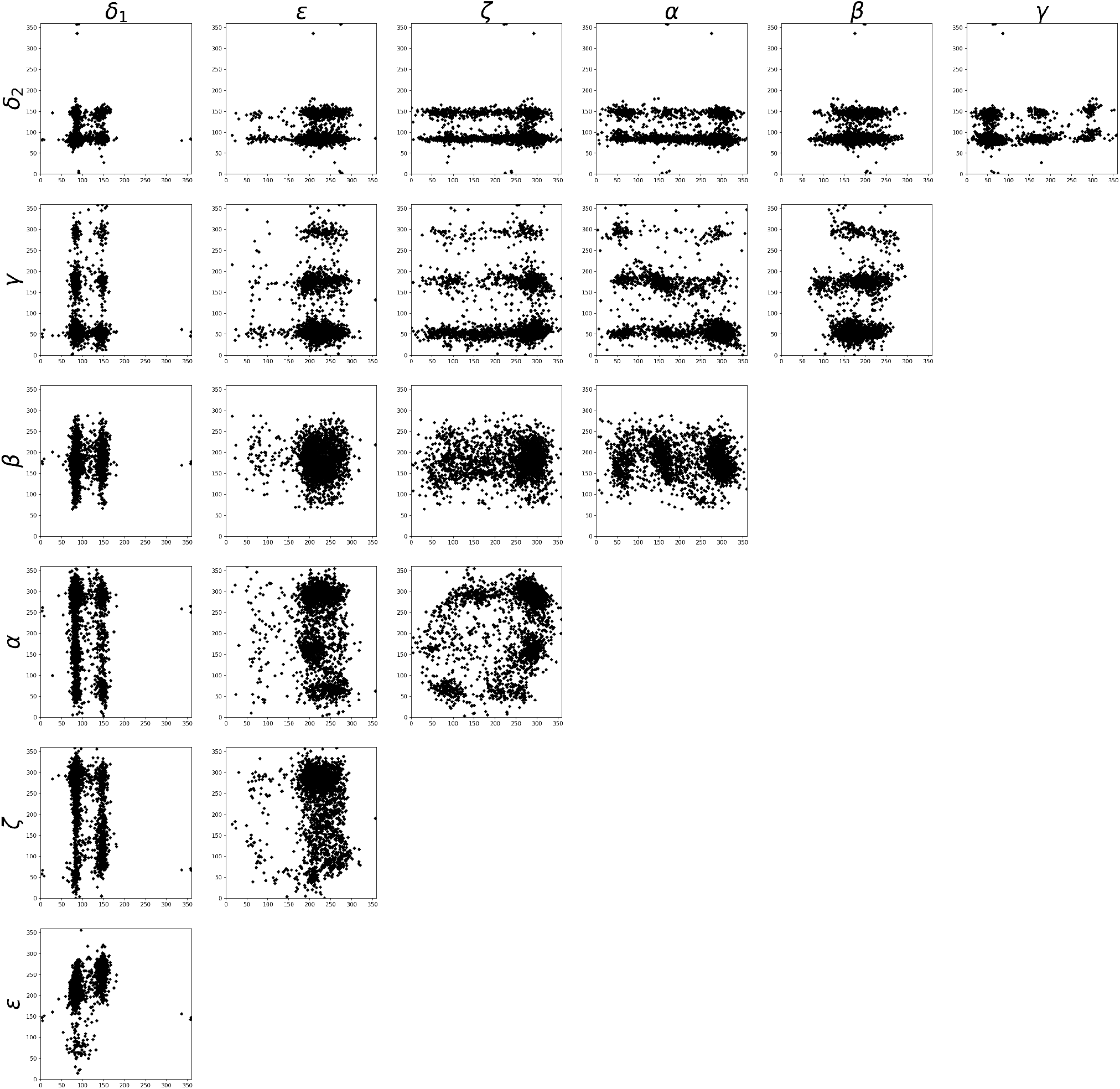
Scatterplots of all two dimensional dihedral angle combinations of the suites of the training data set 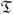, see Supplement D.4.

**Fig. 13:**
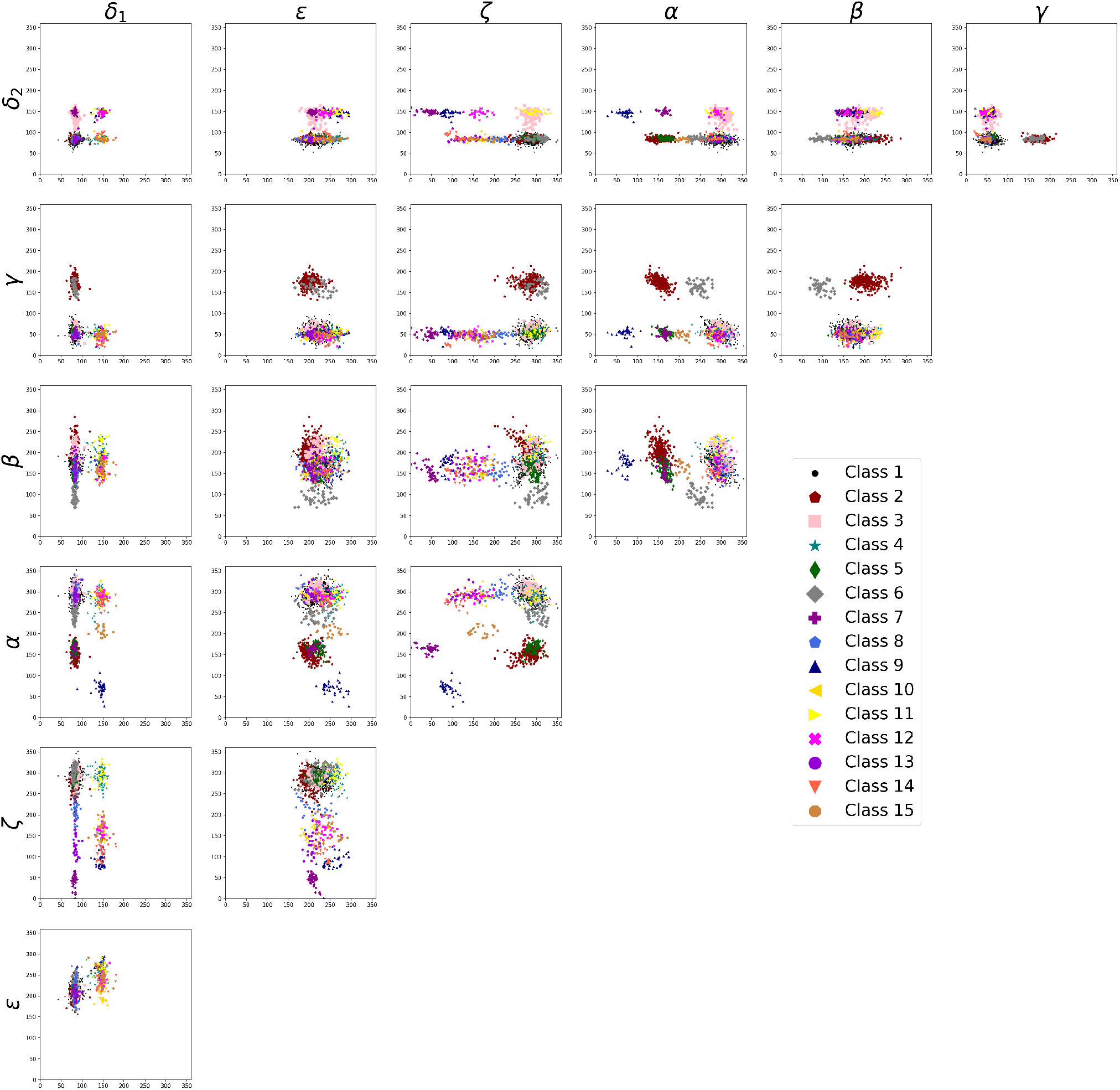
Scatterplots of all two dimensional dihedral angle combinations only for the 15 MINTAGE benchmark classes described in Supplement G.2.

**Fig. 14:**
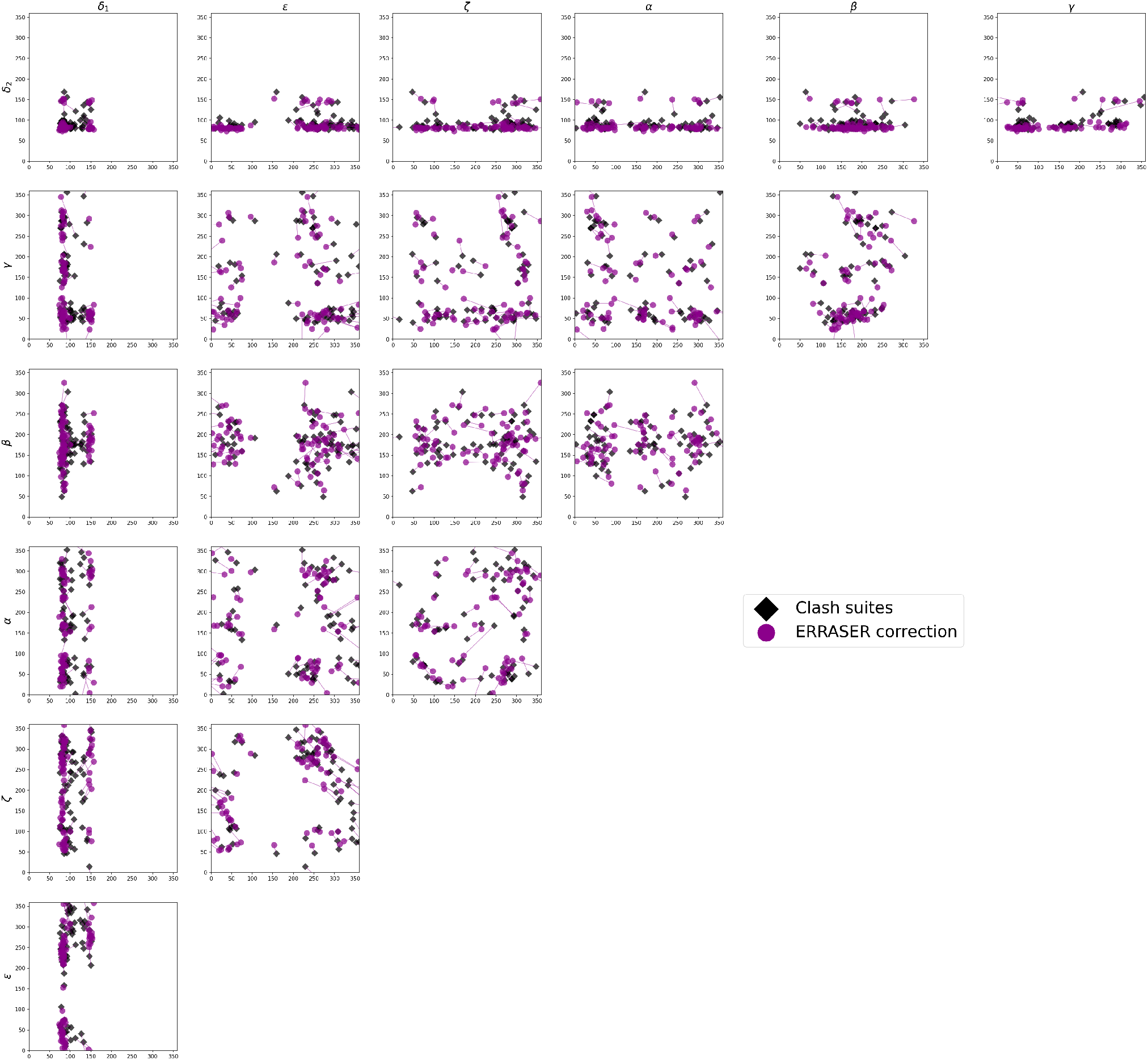
Scatterplots of the 73 clash suites 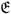 (black) from the benchmark data set with their corrections (magenta) by ERRASER from Chou *et al*. (2013). More than half of them still feature clashes, cf. Supplement I.1.

**Fig. 15:**
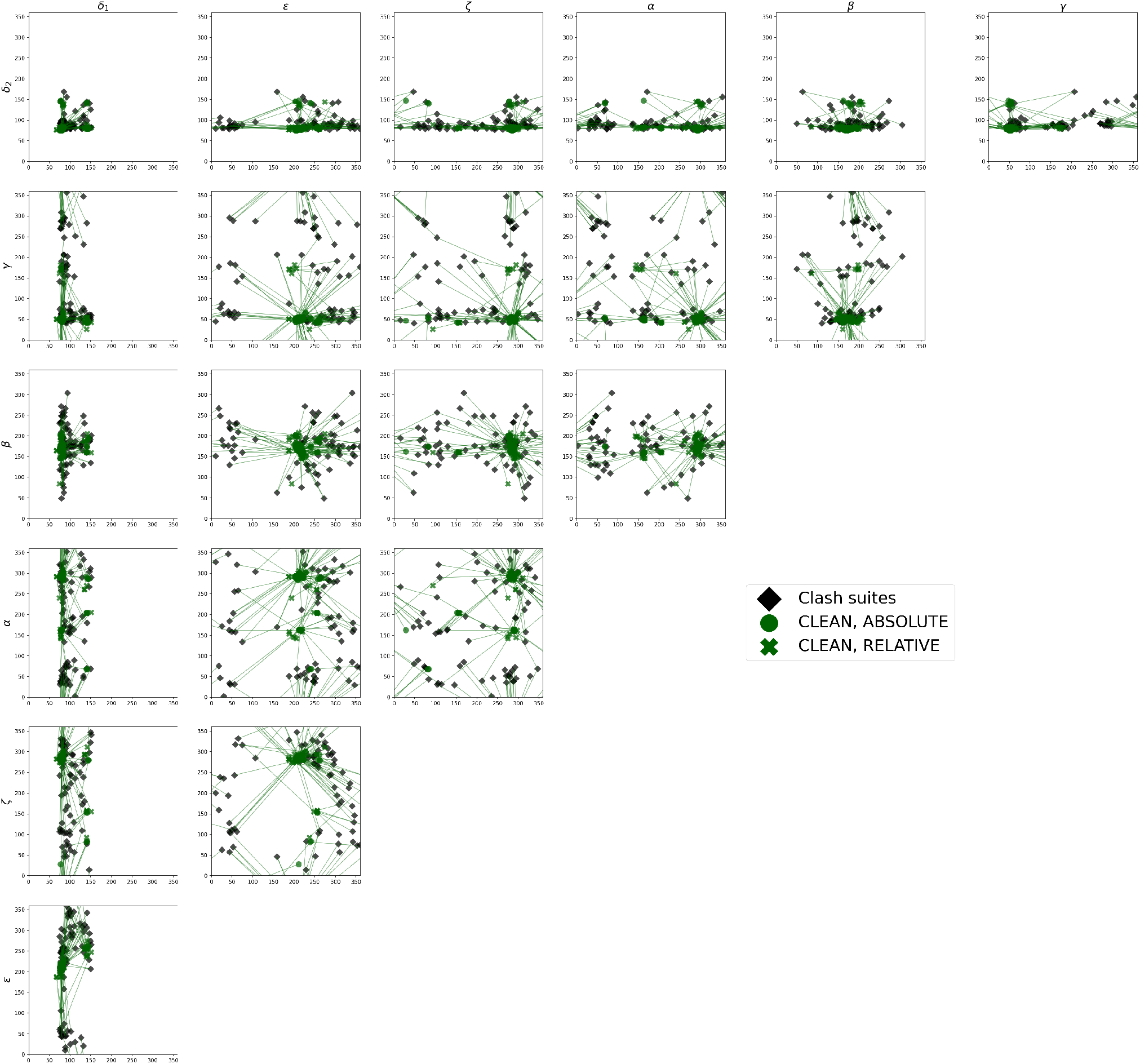
Scatterplots of the 73 clash suites E (black) from the benchmark data set with their corrections (green) by CLEAN MINTAGE (circles correspond to flag DOMINATING = ABSOLUTE in Algorithm 2.4 in the main text and crosses to flag DOMINATING = RELATIVE). Each correction belongs to a clash free class.

### C Fréchet Means and Size-And-Shape Space

We describe geometrical objects by an array of seven dihedral angles on the *torus* 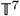 as detailed in Supplement D.2 with the torus distance 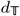 from (4), and by the *size-and-shape* of a configuration matrix 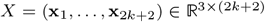 encoding 2*k* + 2, 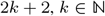, three-dimensional landmark positions 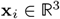, *i* = 1,…, 2*k* + 2 as follows, see Dryden and Mardia (2016).

#### C.1 Size-And-Shape Space

Proper Euclidean transformations comprising rotations and translations 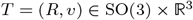 act on *X* columnwise via

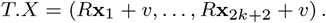

Then

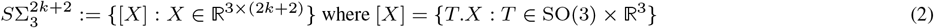

is the *size-and-shape space* which is equipped with the quotient distance, also called *Procrustes distance*

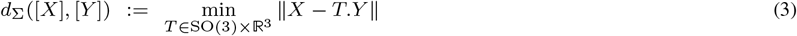

with the standard Frobenius norm on 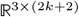. We say that *X* and *Y* are in *optimal position* if

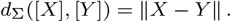

Taking derivatives one can show that configurations *X*, *Y* in optimal position have coinciding mean landmarks with symmetric *XY^T^* (e.g. Dryden and Mardia (2016, Result 7.1)). For this reason, we assume that all landmark configurations are *centered*, i.e. their landmarks vectors add up to zero. Optimal positioning is then conveyed by rotations *R* ∈ SO(3) only.

#### C.2 Fréchet Means

For data *X*_1_,…, *X_n_* ∈ *M* on an arbitrary metric space (*M, d*), define their *Fréchet mean* by

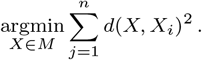

This is a generalization of the classical Euclidean mean. On complete spaces, Fréchet means exist, and on manifolds, if samples are drawn from continuous distributions, they are almost surely unique (see Arnaudon and Miclo (2014)).

On 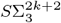, Fréchet means defined by Procrustes distance are also called *Procrustes means*. On 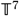 we call them *torus means*.

### D RNA Backbone Geometry and Data Sets

RNA molecules are composed of repeating elements called *nucleotides* and each nucleotide is composed of three building blocks, see Watson *et al*. (2004), cf. Figure 1 in the main text: A sugar ring called *ribose* comprising 5 carbon atoms, one of 4 possible *nucleobases* which is attached to the ribose at the C1’ position and a *phosphate group* connected to the ribose ring at the O5’ atom. The single nucleotides are connected by their O3’ atoms to the next phosphate group to form long RNA chains.

#### D.1 RNA folding

In contrast to DNA which usually forms a double helix (its most common form is B-DNA) of complementary strands, in principle, RNA is single stranded and the form of its ribose (which is not “desoxy” as in DNA, i.e. it has an additional hydroxyl group) allows for complex folding structures. Figure 16 shows a typical RNA structure element, called a *hairpin*. It is composed of double helices (its most common form A-RNA is similar to A-DNA and similarly to DNA, the binding occurs along complementary nucleobases), also called *stems* that are followed by non binding regions called *bulges* and terminated by a terminating region called a *loop*, see for example Wadley *et al*. (2007) and Richardson *et al*. (2008).

**Fig. 16:**
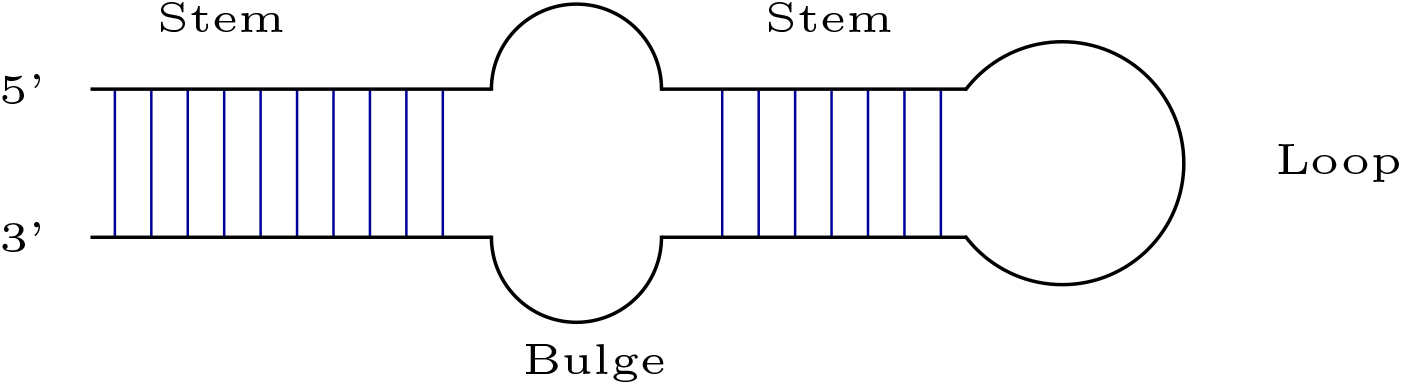
The *hairpin* is a common RNA structure element. Double helices *(stems)* matched by base bindings between nucleotides denoted blue are followed by *mismatching* sites *(bulges)* with several nucleotides not featuring base bindings and a terminating mismatching site *(loop)*. Orientation is conveyed by the 5’ and 3’ ends, cf. Figure 1 of the main text.

#### D.2 Modeling RNA Suites on the Torus

Nucleotides are either studied as *suites*, i.e. from one sugar to the next, or as *residues*, i.e. from one phosphate to the next, e.g. Murray *et al*. (2003); Jain *et al*. (2015). As clashes often occur between neighboring residues but within same suites, cf. Murray *et al*. (2003), for our analysis, we use suites. Indexing, however, is usually done on residue level, so that within a single suite, atom indices change, cf. Figure 1 of the main text. Every directed bond between two atoms (C4’ and C3’, say, cf. Figure 1) defines a *dihedral angle* in the onedimensional *torus*

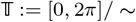

where “~” denotes that 0 and 2π are identified. It is the directed angle between the plane spanned by the bond and the bond preceding (C5’ and C4’, say) and the plane spanned by the bond and the bond succeeding (C3’ and O3’, say), see right panel of Figure 17. For the dihedral angles of concern, Figure 17 lists the 4 consecutive atoms, defining the respective dihedral angles of the bond between the two central atoms.

**Fig. 17:**
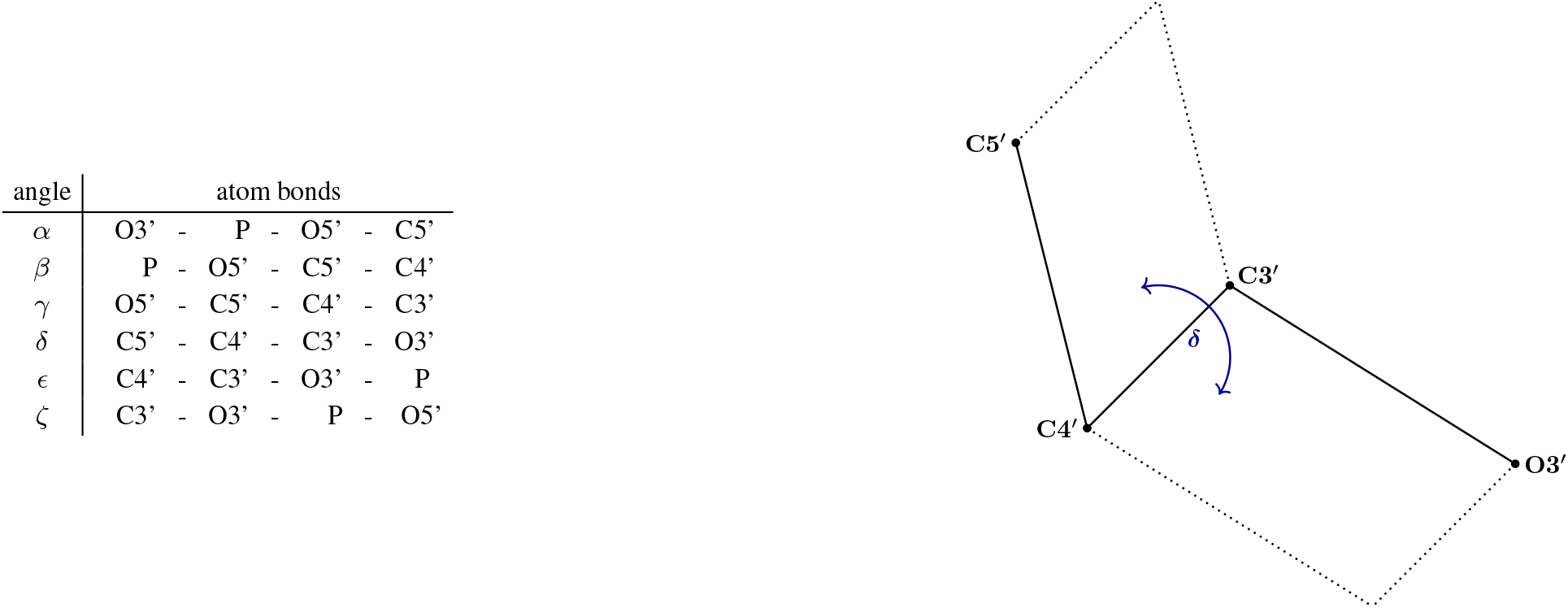
*Left*: Names (first column) of dihedral angles along the two central atoms of the four atoms involved (second column), see Figure 1 in the main text. Right: The dihedral angle δ determined by the atoms C5′, C4′, C3′, O3′.

##### Definition D.1.

*We consider a connected RNA strand with* 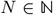 *consecutive nucleotides indexed by i* ∈ {1,…, *N*}. *The i-th* suite *comprises the RNA region between a* 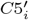 *atom and the second next O*3′ *atom labeled* 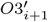 *and the* backbone shape *of the suite is described by the seven dihedral angles* 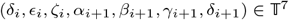 *for i* = 1, …, *N* – *1, cf. Figure 1 in the main text*.

Indeed, geometric suite variability is solely governed by the dihedral angles, since bond lengths (distances between two consecutive atoms) and bond angles (angles between three consecutive atoms) are approximately constant due to the laws of chemistry, Watson *et al*. (2004). In consequence, the geometry of the *i*-th suite is described, up to a proper Euclidean transformation (translation and rotation), by (*δ_i_*, *∊_i_*, *ζ_i_*, *α*_*i* + 1_, *β*_*i*+1_, *γ*_*i*+1_, *δ*_*i*+1_) on the seven-dimensional torus. 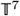 is a metric space with the canonical product distance 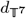 where

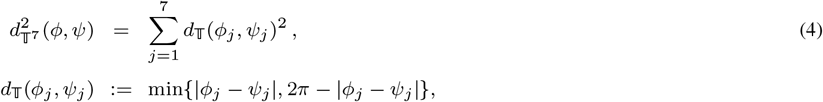

with *j* = 1,…, 7 and 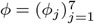, 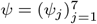.

#### D.3 Size-And-Shapes of Mesoscopic Strands

The mesoscopic view takes the coordinates of the 2*k* + 2 ‘closest’ sugar rings into account and can be seen as an intermediate scale between the suite or microscopic (single atoms) and macroscopic (the whole RNA strand) representation.

#### Definition D.2.

*We consider a connected RNA strand with N* ≥ 2*k* + 2 *consecutive nucleotides indexed in i* ∈{1,…, *N*}, *see Figure 4 in the main text. Each nucleotide comes with a sugar ring formed by the atoms* 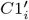, 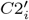, 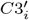, 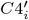 *and* 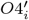 *(see Figure 1 in the main text). Denoting their center of gravity (i.e. average location) with* 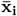, *for all i* = *k* + 1,…, *N* – *k* − 1, *the* mesoscopic strand *corresponding to the i-th suite is the* configuration matrix 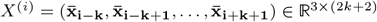. *Its size-and-shape in* 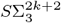 *see Dryden and Mardia (2016), is called its* mesoscopic shape.

Since distances between two neighboring sugar rings and angles between three consecutive sugar rings vary, dihedral angles defined by four consecutive sugar rings are not sufficient to completely define the geometry of mesoscopic strands up to proper Euclidean transformations. The size-and-shape representation, however suffices.

#### Remark D.3.

*For the mesoscopic strands we include the sugar ring centers of the k* = 2 *suites preceding and the k* = 2 *suites following the suite of concern, cf. Figure 4 in the main text. Choosing k* = 2 *reflects the local geometry at an intermediate (mesoscopic) scale as the* 5 + 1 = 6 *bases from the* 2*k* + 1 = 5 *suites correspond roughly to the number of bases involved in a half helix turn, see e.g. *Watson* et al. *(2004)**.

We only work with suites that have a corresponding mesoscopic strand, i.e. suites that are not on the outer edge of an RNA strand.

#### Definition D.4.

*For an RNA strand of length N* ≥ 2*k* + 2 *the suites numbered i* = *k* + 1,…, *N* – *k* – 1 *are called* admissible, *so that every admissible suite a has a mesoscopic shape* 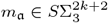 *and vice versa*.

### D.4 The Benchmark, Training and Test Data Sets

In our applications, we analyze a subset of a classical RNA data set. The classical RNA data set comprises 8665 suites, carefully selected for high experimental X-ray precision (of 3 Å = 0.3 nanometers) by Duarte and Pyle (1998); Wadley *et al*. (2007) and analyzed by them and by others, for example Murray *et al*. (2003); Richardson *et al*. (2008) and Eltzner *et al*. (2018). The data originate from 71 different measurements and the atomic positions of each measurement have been stored in the *PDB* format of a *protein data bank* file, online at the Protein Data Bank, see Berman *et al*. (2000). More details on the PDB files can be found in Table 1 of Supplement A.

From this classical data set, we consider the 7648 admissible suites (which have an associated mesoscopic strand, see Definition D.4) and call this data set the *benchmark data set*.

- From the benchmark data set the 5957 clash free suites (suites that also have clash free mesoscopic strands, see Definition 1.2 in the main text) form the *benchmark training data set* 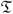, see Figure 12.
- From the remaining suites we chose the suites that feature within suite clashes, forming the *benchmark test data set* 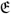, containing 198 suites.

As our purpose lies in demonstrating our methods rather than correcting all clashes, all other suites (not themselves clashing but featuring clashes in their mesoscopic strands) are disregarded in our analysis.

### E Cryo-EM, X-Ray Crystallography, Validation and Correction

Cryo-EM (cryogenic electron microscopy) and X-ray crystallography are popular methods to determine atomic positions in RNA structures, cf. Jain *et al*. (2015). For the former, RNA molecules are shock frosted and subjected to electron microscopy. For the latter, using a suitable substrate, RNA molecules are crystallized and subjected to X-ray imaging. The resolution of X-ray crystallography is defined as the smallest distance of two objects such that their diffraction patterns can be separated. In cryo-EM, resolution has been defined in various ways, usually via properties of the Fourier transformed electron density, in order to be comparable to the resolution values given for X-ray crystallography measurements. For a review, see Liao and Frank (2010). From different angles, via inverse Fourier transforms, the electron density can be reconstructed and, in principle, density peaks correspond to atom positions. Figure 18 shows exemplary level surfaces of electron densities with estimated atom positions.

**Fig. 18:**
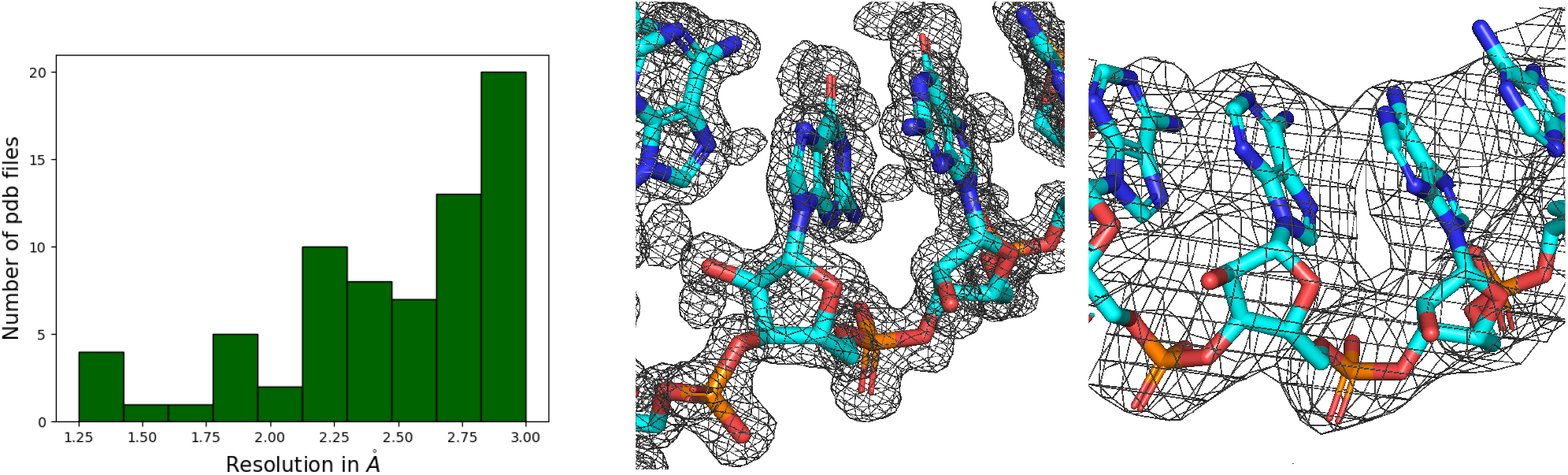
Left: Histogram of X-ray crystallography resolutions in the benchmark data set, see Supplement D.4. Middle and right: reconstructed RNA structure and electron density contour surface created with PyMOL at level of one σ, see Schrödinger, LLC (2015), at resolution 1.6 Å (middle, from the benchmark file 1csl, Ippolito and Steitz (2000), see Table 1 building our benchmark data set in Supplement D.4) and at resolution 3 Å (right, from benchmark file 1f8v, Tang *et al*. (2001)).

At a resolution of 2.5 to 4*Å*, which is typical for large RNA strands, base pairings can be predicted well and phosphates are well identified by strong peaks of density Jain *et al*. (2015). It is, however, more challenging to precisely estimate the atom positions describing the sugar-phosphate backbone, see for example Murray *et al*. (2003). In addition, structural disorder due to crystallization and thermal oscillation contribute to uncertainties, occasionally resulting in incompatible reconstructed atom positions. Indeed, our benchmark data set contains approximately 2.5 % clash suites.

The PHENIX (Python-based Hierarchical ENvironment for Integrated Xtallography) software provides some validation tools that detect local errors, see Liebschner *et al*. (2019). While there are different types of errors detected by PHENIX, in this work we focus on backbone clashes, see Definition 1.2 in the main text. They usually occur after the hydrogen atoms (cf. Figure 1 in the main text) which are not visible in the electron density measurements (H-atoms contain only one electron which is shifted to the covalent-bond partner atom) are added to the reconstructed configuration.

A popular and well established (see for example Richardson *et al*. (2018), Jain *et al*. (2015)) tool for structure correction is ERRASER (Enumerative Real-space Refinement ASsisted by Electron-density under Rosetta) Chou *et al*. (2013). In contrast to manual correction tools such as the RNA Rotator Tool which is available as KiNG, see Chen *et al*. (2009), ERRASER automatically corrects complete RNA structures as well as individual residues. It usually executes the following three steps three times in succession, see Chou *et al*. (2013):

1. The high-resolution *Rosetta energy function*, extended by *electron density correlation evaluation*, subjects, among others, all dihedral angles to minimization in order to obtain a new reconstruction.
2. Residues in this new reconstruction are labeled if the PHENIX validation tools (Supplement F.1) detect errors, if the backbone’s configuration is not recognized, or if other geometric errors occur (e.g. in the structure of the sugar ring).
3. Labeled residues are reconstructed one after the other by Single Nucleotide StepWise Assembly (SWA) which samples all nucleotide atoms from an exhaustive grid search.

Supplement F details how we use and apply PHENIX and ERRASER. In application to the benchmark data set from Supplement D.4 we compare ERRASER to our proposed CLEAN MINTAGE in Section 3.1 of the main text and in Supplement I.

## F Using PHENIX and ERRASER

### F.1 PHENIX

Since hydrogen atoms are not visible in the electron density measurements, first, the PHENIX tool `phenix.reduce` adds the hydrogen atoms. Then, `phenix.probe` performs an all-atom contact analysis (Word *et al*., 1999), which declares atoms that are not bonded to each other as a *clash* if they are closer together than is physically possible (i.e. if van der Waals shells overlap by more than 0.4 Å). For each PDB file, `phenix.clashscore` generates a list of all clashes.

### F.2 ERRASER

To correct a PDB file with ERRASER, one needs a 2mFo-DFc density map in CCP4 format in addition to the raw PDB file. We created the 2mFo-DFc electron density maps with the PHENIX map tool, see Liebschner *et al*. (2019). However, some older PDB files have not published the associated experimental files necessary to create a 2mFo-DFc density map in CCP4 format and were therefore left out. We used the offered online server ROSIE (Rosetta Online Server that Includes Everyone) see Chou *et al*. (2013) to obtain the ERRASER corrections. ERRASER returns a statistic for each corrected PDB file that identifies a *clashscore* (the number of clashes per 1000 atoms) in the raw data set and the clashscore in the PDB file corrected by ERRASER.

## G MINTAGE Details

### G.1 Adaptive Cutting Average Linkage Tree Clustering on a Metric Space

The *average linkage clustering*, also known as *unweighted pair group method with arithmetic mean* or simply as *UPGMA*, was first developed by Sokal and Michener (1958). It is a hierarchical clustering method that creates a rooted tree where each node stands for a cluster comprising all leaves below that node.

Starting with data *X*^(1)^,…, *X*^(*n*)^ in a metric space with distance *d*, initially, each *X*^(*i*)^ (*i* = 1,…, *n*) is assigned its own cluster yielding the initial *running cluster list*. The tree constructed starts with a graph comprising *n* leaves labeled from 1 to *n*, representing each of these initial clusters. Then, iteratively, if there is more than one cluster in the running cluster list, the two clusters with the smallest average distance are merged to form a new cluster which is added to the running list, from which the two merged clusters are deleted. Here, the *average distance* between two clusters *A* and *B* is given by

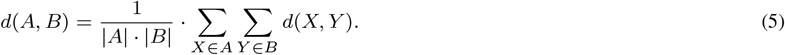

The graph is extended by adding a parent node above the two nodes corresponding to the merged clusters and adding branches joining them to the parent node. If there have been *m* nodes, the parent node is labeled *m* + 1 and assigned a value denoting the distance between the two merged clusters. The iteration terminates if the running list contains only one cluster, comprising all data, which forms the root of the tree, called the *average linkage clustering tree*. Notably, node values increase when approaching the root. Indeed, if clusters A and B are merged to cluster *A* ∪ *B* and *C* is any other cluster, then *d*(*A, B*) ≤ *d*(*A, C*), *d*(*B, C*) and hence

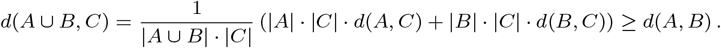

Every choice of a distance value *d_c_* > 0 yields a clustering by a *tree cut* at distance *d_c_*, i.e. by taking that running list when the last node with distance value ≤ *d_c_* has been added. A simple example is illustrated in Figure 19.

**Fig. 19:**
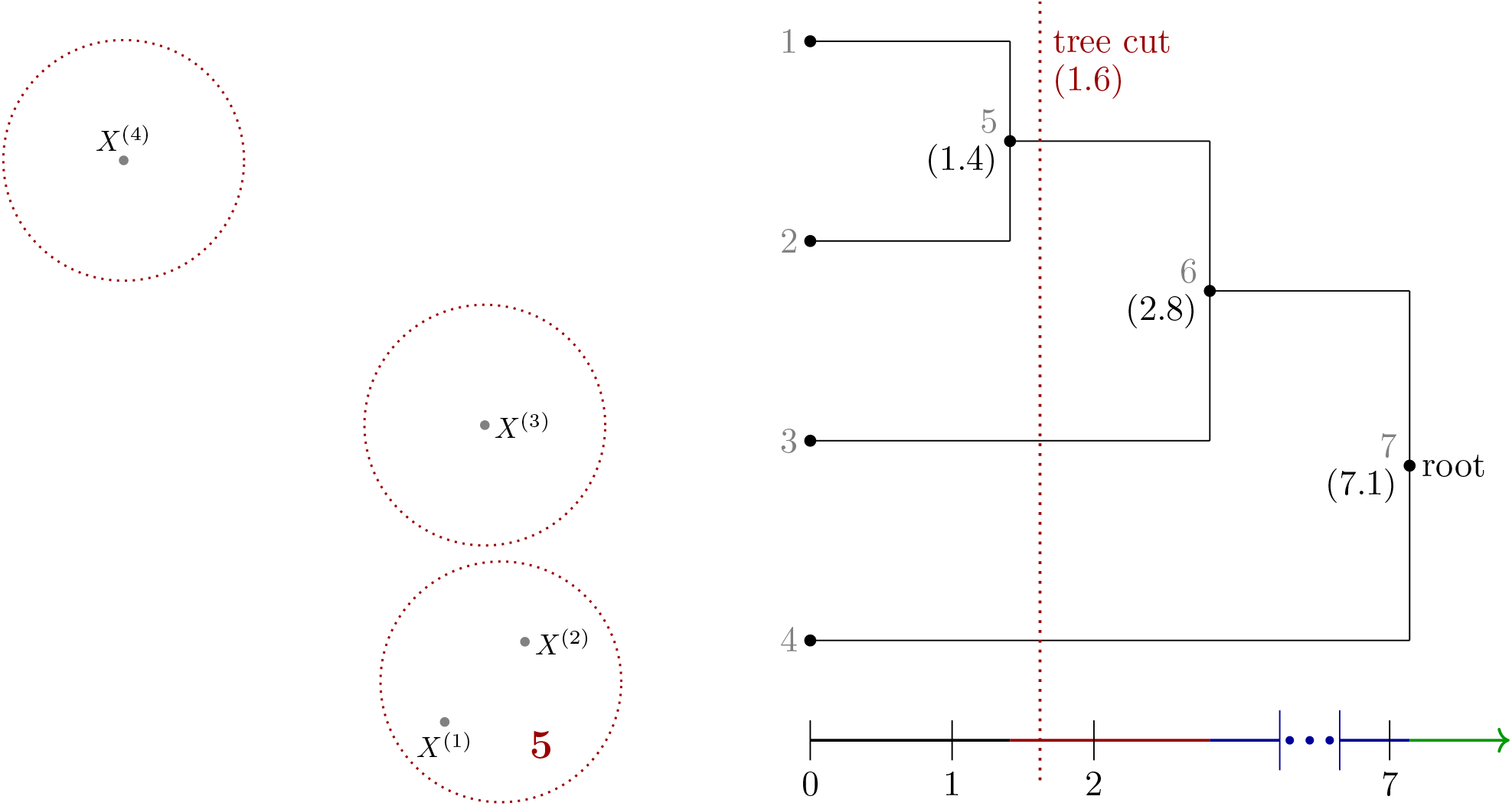
Left: Four data points *X*^(1)^,…, *X*^(4)^ in the Euclidean plane. The dark red circles show the running cluster list after the first merge. Right: The corresponding average linkage cluster tree. The dark red line displays the tree cut at distance 2 (after the first merge). The node values are in parentheses.

We note that instead of the average distance function in (5), one may consider the minimal cluster distance yielding *single linkage* or *nearest neighbor clustering*, developed by Florek *et al*. (1951), which is currently also highly popular. It tends to return long elongated clusters, an effect called *chaining*, see Everitt (1993). For our purpose of structure correction relying on Fréchet means, which are generalizations of averages to metric spaces, clustering based on average distance ensures that Fréchet means are within or close to their clusters with a tendency towards isotropic spread.

In the *simple clustering approach*, the cluster tree is cut at some fixed distance, cf. Figure 19. We use this for motivating our CLEAN algorithm in Section 2.3.1. Obviously, this simple approach fails to separate frequent cluster configurations, for instance if two closely neighboring clusters have a higher density than a third one. Such configurations are separated, however, using a data-adaptive cutting procedure, see for example Langfelder *et al*. (2007) and Obulkasim *et al*. (2015). We choose the following tuning parameters:

- *maximal outlier distance d*_max_ controlling cluster density,
- *minimal cluster size κ* controlling, if not too small, that our mode hunting from Section 2.2 in the main text can significantly separate one-dimensional clusters with several modes,
- *relative branching distance q* ensuring that two clusters are only split if their parent node’s distance value is significant in relation to the greatest distance value of its child nodes.

#### Algorithm G.1.

Consider the set *P* = {*X*^(1)^, …, *X*^(*n*)^} of *n* data points. Let *R* be the outlier list and 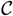 be the cluster list, each of which are initially empty. They are filled iteratively as follows:

1. Compute the average linkage clustering tree from *P*
2. Perform a tree cut at distance *d*_max_ to obtain a clustering, move from *P* to *R* all data points that are in clusters with less than *κ* data points.
3. Compute the average linkage cluster tree for the new *P* as in Step 1.
4. Set 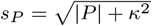 (inspired by the square root rule of thumb used in histogram binning), create an empty list 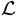 of clusters.
5. Begin at the root and always follow the branch with more points at each node. From each node add the child node corresponding to the smaller subcluster to 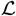

a. if it contains more than *s_P_* data points,
b. and if the *q*-fold of its parent node’s distance value is greater than the two children nodes’ distance values.
6. At the last node, where the smaller subcluster is added to 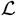, also add the larger subcluster to 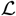
7. Consider 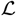:

- if 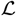 is empty, move the union of all data points from *P* to 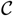; these correspond then to one single cluster,
- else, add the largest cluster in 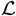 to 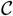 and remove its points from *P*.
8. If |*P*| > 0, go to Step 1.
9. Return the clusters list C and the outlier list *R*.

Figure 3 in the main text illustrates that some of the pre-clusters resulting from the benchmark training set obviously contain subclusters that have not be identified by Algorithm G.1. It turns out they can be well separated by mode hunting (see Section 2.2 in the main text) applied to their one dimensional torus PCA representation.

#### G.2 The 15 MINTAGE Benchmark Classes and the MINTAGE Benchmark Outliers

In this section we cluster the training set suites from Supplement D.4 with the MINTAGE Algorithm 2.1 yielding ***the 15 MINTAGE benchmark classes*** and ***the MINTAGE benchmark outliers***. Since we use this clustering in Section 2.3 in the main text to suggest corrections for clash suites, we aim at larger and concentrated clusters, possibly at the price of a larger number of outliers which cannot be allocated to any of the clusters. To this end, for the pre-clustering Algorithm G.1 we choose the *minimal cluster size κ* = 20, the *relative branching distance q* = 0.15 and the *maximal outlier distance d*_max_ such that 5% of the suites in the average linkage tree are in a branch with less than *κ* data points. The clustering results are summarized in Figure 20.

**Fig. 20:**
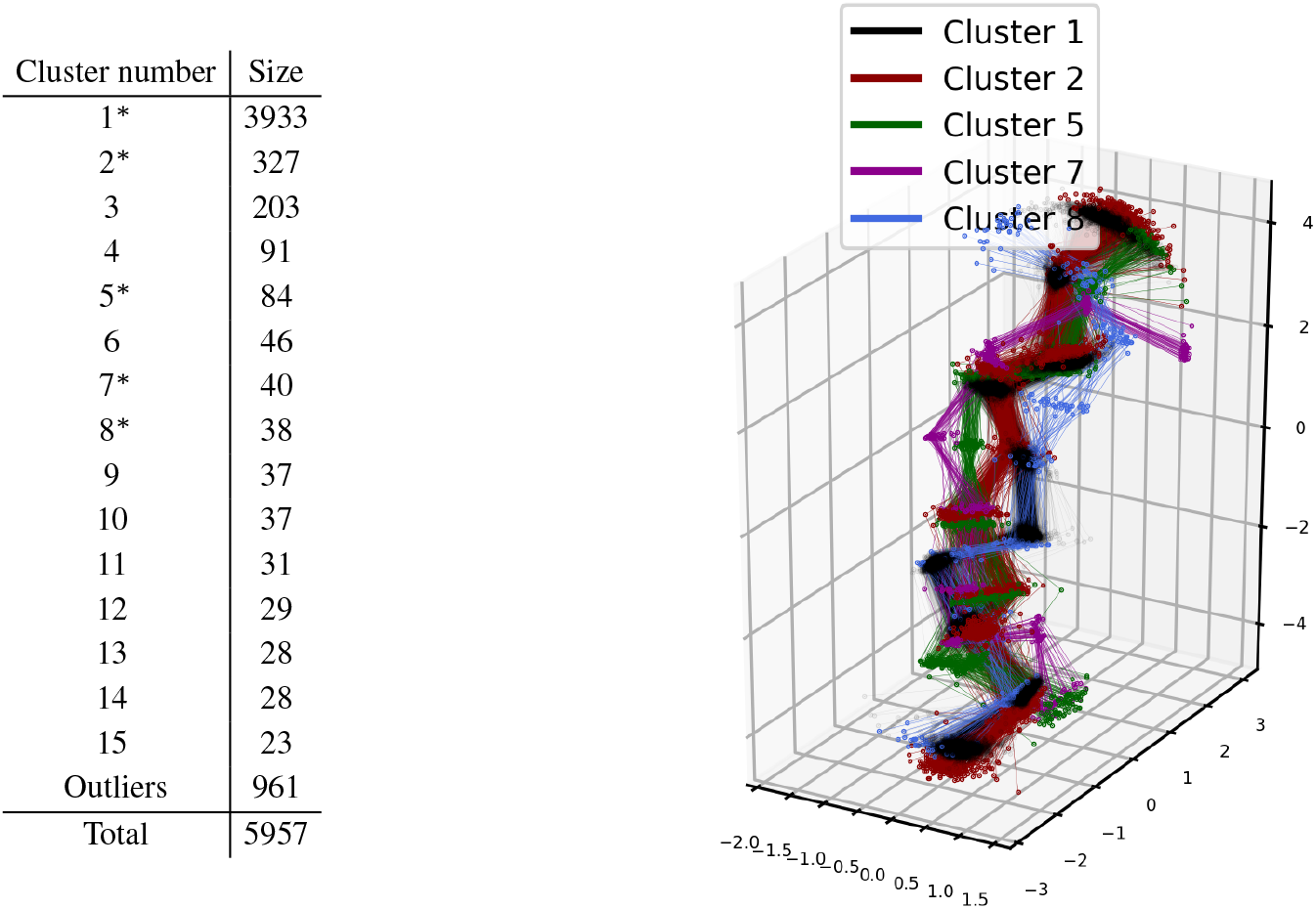
Left: Clusters and outliers (numbers and sizes) obtained from applying MINTAGE to the benchmark training data set yielding **the 15 MINTAGE benchmark classes and outliers**. Asterisks mark clusters displayed in the right panel. Right: Five exemplary clusters that can be well displayed together (colors conforming with Figure 3 in the main text).

With the tuning parameters thus set, our torus PCA based MINTAGE method returns 15 clusters. The largest corresponding to the A helix shape contains 3933 elements and is highly dominant. All clusters are rather dense and even the smallest cluster has a credible size of 21 elements. The number of outliers (961), however, is quite large. We conjecture that a considerable number of these are due to incorrect structure reconstructions, which have not been detected because they have not led to clashes. All 15 clusters determined with the MINTAGE clustering are also displayed in Figure 13.

### H CLEAN Details

#### H.1 Motivation, Procrustes Step and Design

The following Figures 21 and 22 complement the motivation of CLEAN in Section 2.3.1 from the main text.

**Fig. 21:**
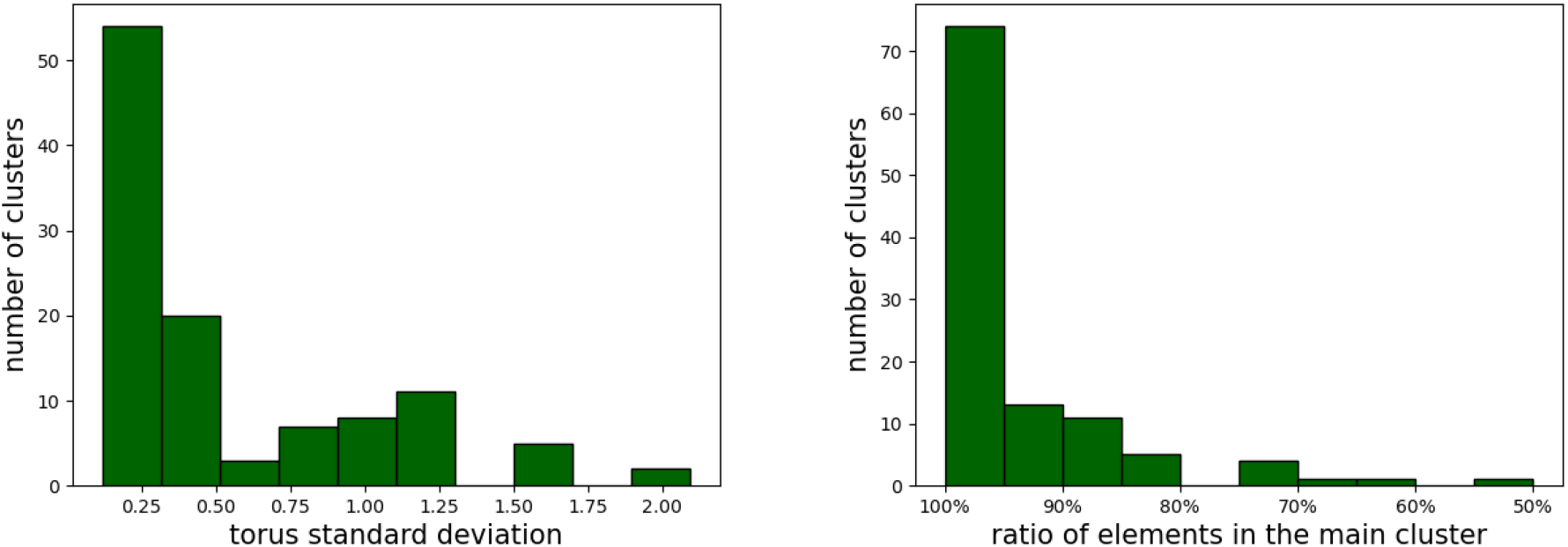
Left: Binned torus standard deviation of the suites belonging to simple mesoscopic clusters. For instance, the suites of Cluster 1 from Figure 7 in the main text have a standard deviation of 0.83, so that Cluster 1 is counted in the 4th green bar from the left. Right: Percentages of suites in simple mesoscopic clusters belonging to a single MINTAGE benchmark cluster.

**Fig. 22:**
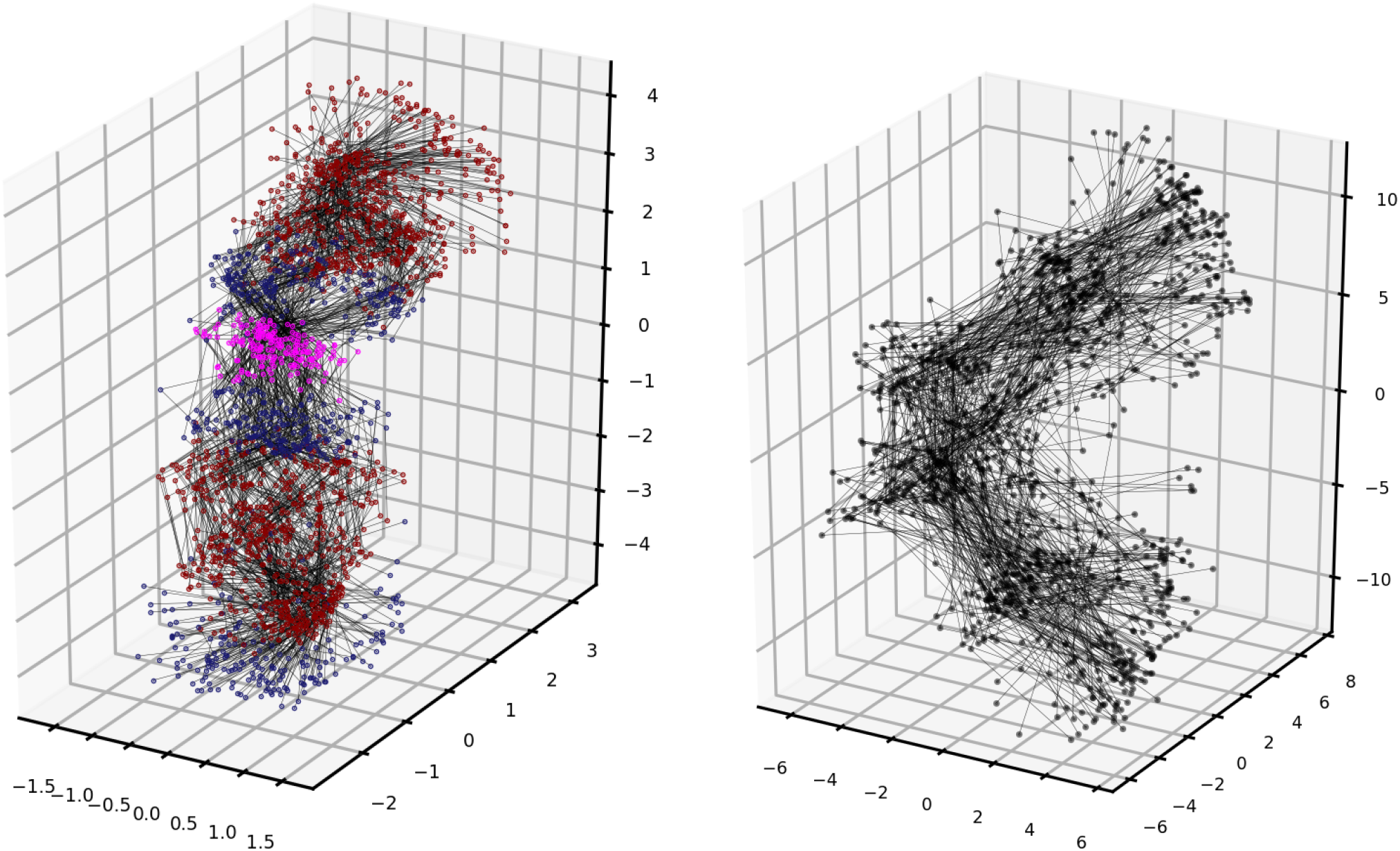
Left: The clash suites from the benchmark data set in Supplement D.4 with carbon (dark red), oxygen (dark blue) and phosphorus atoms (pink), cf. Figure 1 in the main text. Right: Their mesoscopic shapes.

##### Algorithm H.1.

The Procrustes Step 2 in the CLEAN Algorithm 2.4 in the main text, with notation given there, proceeds as follows:

1. With 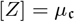 and 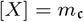 where *Z* = (*z*_1_,…, *z*_2*k*+2_) and *X* = (*x*_1_,…, *x*_2*k*+2_), set (*y*_1_,…, *y*_2*k*+2_) = *Y* ≔ *Z*
2. Obtain *Y*′ from *Y* by setting *Y*′ ≔ *Y* and additionally

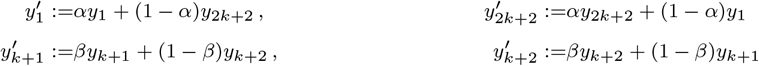

where

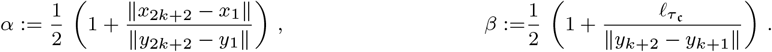
3. Rotate *Y*′ into optimal position 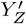 with respect to *Z* and set 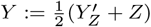
4. Repeat Steps 2 and 3 until [*Y*′] converges.

##### Remark H.2.

*By design, each suite corrected by the CLEAN Algorithm 2.4 is assigned to one of the torus PCA clusters. Moreover, we have:*

1. *The algorithm uses the ρ most similar mesoscopic shapes* 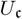. *Throughout the applications, we chooses ρ* = 50 *which corresponds to about one percent of the benchmark training data. Further, choosing a locally dominating and not a globally closest cluster, we prevent the algorithm from displacing landmarks of the mesoscopic shapes too drastically*.
2. *As scale is not factored out in size-and-shape space, in Step 3 of Algorithm 2.4 the distance between Landmarks 3 and 4 is set to the length of the Fréchet mean suite, to ensure that* 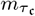 *is indeed a mesoscopic shape compatible with* 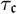. *Also the distance between Landmarks 1 and 6 remains constant. Hence*, 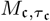 *is a proper subspace of* 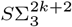 *which will usually not contain the Procrustes mean* 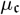.
3. *Since geodesics in* 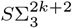 *are projections of straight lines between preshapes in optimal position (e.g. *Huckemann* et al. *(2010)*), the orthogonal projection* 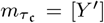 *of* 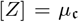 *to* 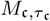 *is a fixed point of the Subalgorithm H.1 of Algorithm 2.4. In applications we observe rather fast convergence*.

#### H.2 Validation of CLEAN

We apply the CLEAN Algorithm 2.4 to the 198 clash suites which form the test data set 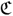, which is part of the benchmark data set from Supplement D.4.

For validation we reassure ourselves that backbone correction is realistic and neither arbitrary nor ambiguous. For the former, verify that corrections happen on a scale not larger than the underlying X-ray crystallography resolution, see Supplement E, and for the latter we verify that the largest clusters in neighborhoods 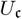 are indeed strongly dominating in most cases.

In order to relate the amount of correction to resolution, consider

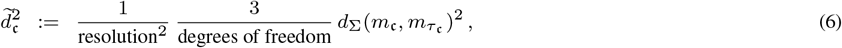

for 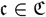. The *degrees of freedom* are given by 3(2*k* + 2) −3 −2 −1 = 3 · 2*k*, so that the inverse of the second quotient above gives the number 2*k* of free landmarks in 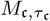.

The histogram in Figure 23 shows that for the vast majority of clash suites 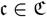, 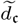 is smaller than 1. Thus, corrections are only rarely slightly above and mostly well below the order of resolution.

**Fig. 23:**
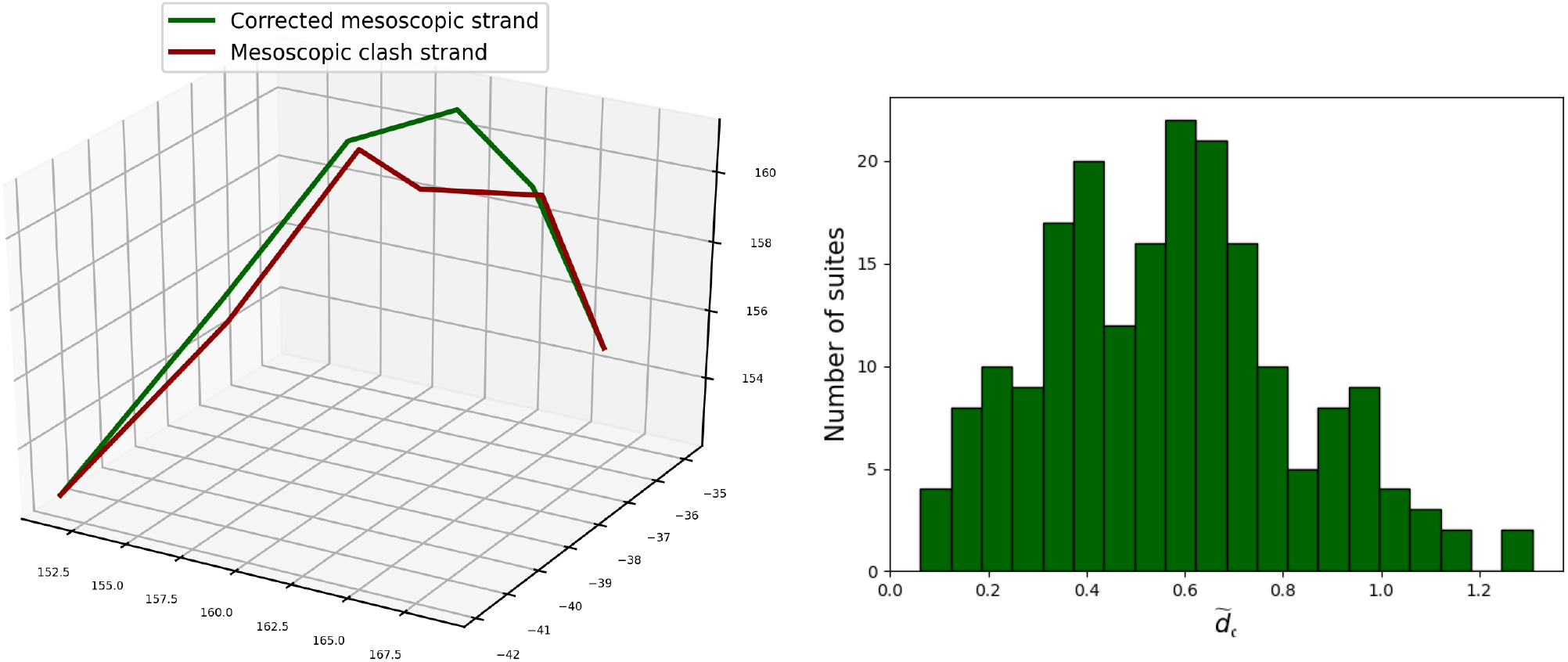
Left: A mesoscopic strand of a clash suite (red) and its corrected mesoscopic strand (with equal first and last landmark) rotated into optimal position (green) of the torus mean of the dominating cluster in 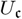 (defined in (1) in the main text, not depicted). Right: Histogram of relative distances 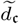, 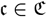, from (6) between corrected mesoscopic shapes and original mesoscopic shapes.

In order to assess how dominating torus PCA clusters are in neighborhoods 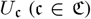, the histogram in Figure 24 shows the number of suites in the dominating clusters 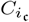. Indeed, for many more than half of the neighborhoods, the dominating cluster contains more than half of the neighboring suites. Remarkably, the negative correlation visible in the scatter plot in Figure 24 (right) shows that a smaller amount of correction tends to correlate with more elements being in the dominating cluster. c

**Fig. 24:**
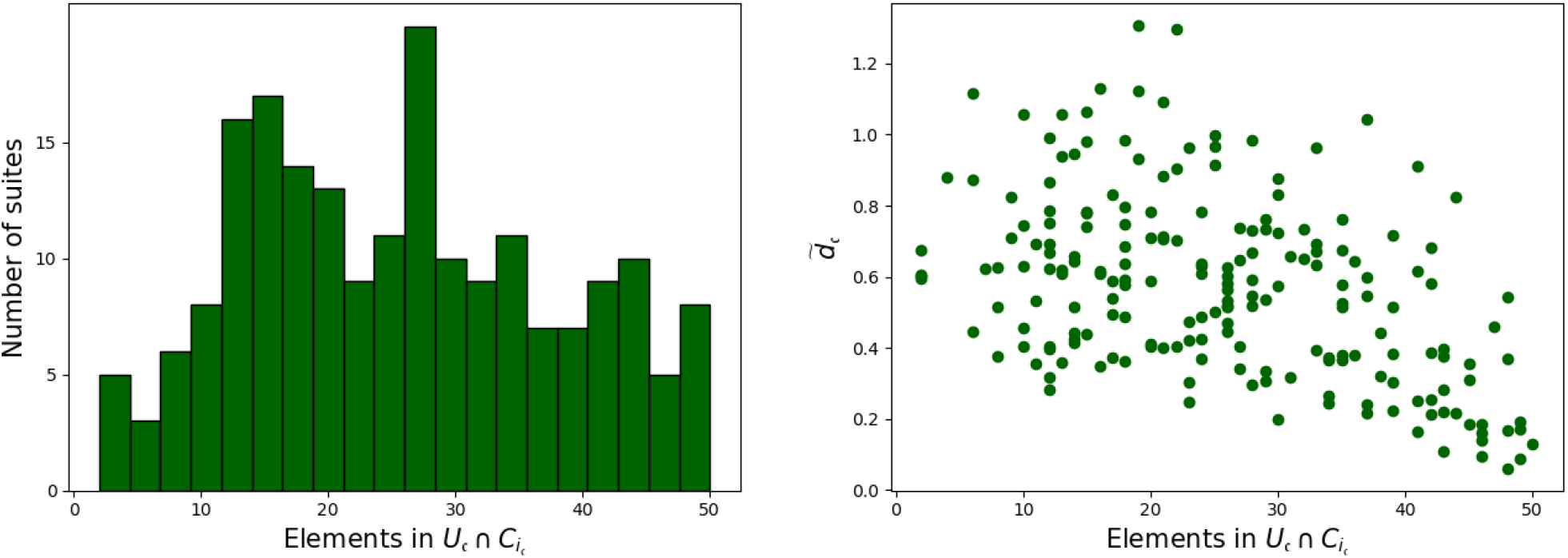
Left: Histogram of the number of suites in 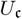 (defined in (1) in the main text), 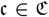, of the dominating torus PCA cluster. Right: Scatter plot relating the number of suites in the dominating cluster of 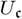 and the relative distance 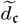, over 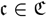.

### I Comparing CLEAN MINTAGE with ERRASER

By design, as mentioned in Remark H.2, our CLEAN MINTAGE algorithm corrects clashes by assigning clash suites to one of the classes of the underlying training data set. In contrast, after correction by ERRASER, see Supplement E, many clash suites remain outliers. We show this in this section by applying MINTAGE to the training data set 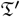 (defined below) which is closely related to the benchmark data set, specifically tailored to ERRASER.

In fact, ERRASER seems to correct rather locally and cautiously and thus only part of the clashes, whereas our algorithm corrects more globally into configurations near the training data, that are thus most likely clash free, see Figures 2 and 9 from the main text and Figure 14.

Notably, the ERRASER method on the ROSIE server can only correct PDB files that come with an associated 2mFo-DFc density map in CCP4 format and do not exceed a specific maximum size. This is the case for only 49 PDB files of the 71 PDB files from our benchmark data set, comprising 2325 suites. We denote this set by 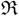 and it is a subset of our benchmark data set from Supplement D.4. Whether clashing or not, all of the suites from 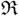 are corrected by ERRASER (processing PDB files) and we denote the corrected data set with 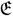, and again, 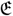 comprises 2325 suites.

As the data set 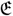 is too small for the learning step of CLEAN MINTAGE, we augment it by the clash free suites 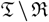 (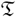 denotes the original training data set from Supplement D.4) which are not corrected by ERRASER, to obtain the

- the *ERRASER training data set* 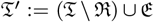, which has now the size 6433 (it comprises slightly more suites than 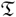, namely also of the former clash suites and its neighbors after correction by ERRASER),
- the *ERRASER test data set* 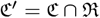 of clash suites which has size 73, and
- he *ERRASER data set* 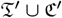 of size 6506.

#### I.1 Clash Reduction by ERRASER

We apply the validation method PHENIX, see Supplement E, which comes with the tool `phenix.clashscore` which returns the number of clashes per 1000 atoms found in a given PDB file. Note that this *clash score* includes all clashes between two atoms in the measurement, not just the clashes between two backbone atoms.

The left panel of Figure 25 shows that ERRASER effectively reduces clash scores as an overwhelming part of the data points (each corresponding to a single PDB file) lies substantially below the diagonal. The right panel of Figure 25 shows the amount of clash reduction on suite level: after correction by ERRASER, from 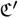, approx. 40 % (29) of the suites have been made clash free and almost equally many (30) feature only one clash; correcting the few suites with a higher number of clashes was only partially possible.

**Fig. 25:**
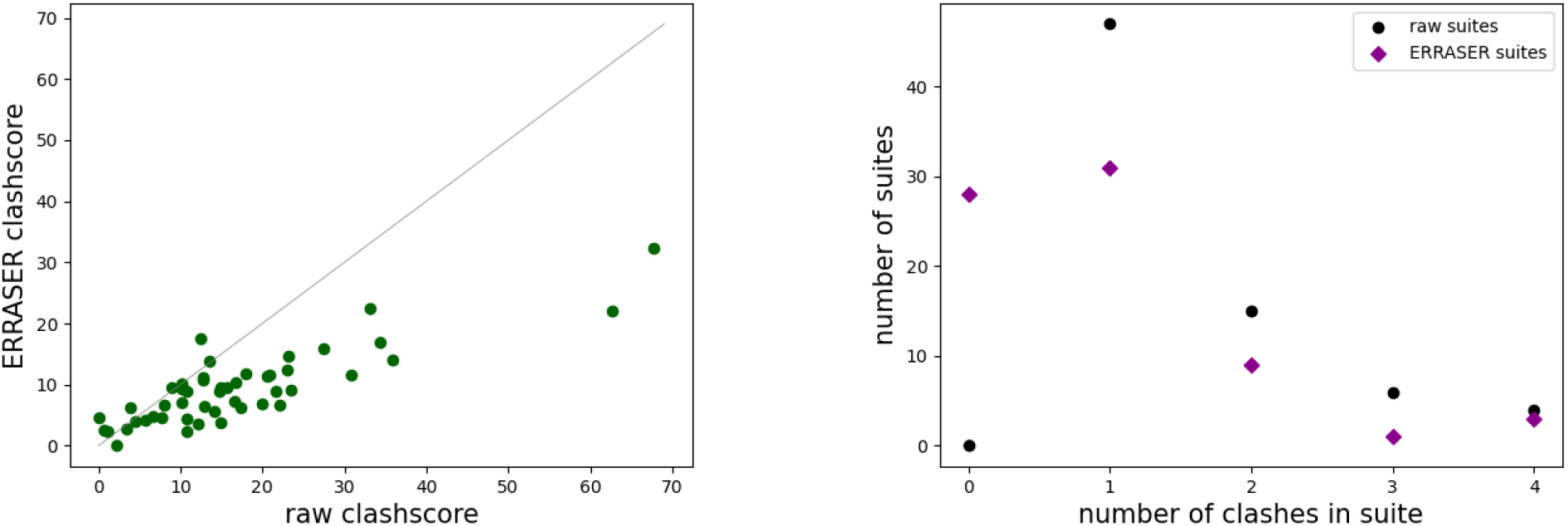
Left: Each of the 73 green points compares the clash score of a single PDB files before (horizontal) and after (vertical) correction by ERRASER. Right: Histogram of numbers of atom clashes (horizontal) in clash suites before (black) and after correction by ERRASER (magenta).

Figure 2 in the main text displays the clash suites from 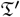 before and after correction by ERRASER and CLEAN MINTAGE. Through ERRASER correction, no distinct structures become visible. As illustrated in Supplement I.2, a majority of suites remain outliers even after correction by ERRASER. In stark contrast, CLEAN MINTAGE assigns most clash suites to the dominating (first cluster, A helix) and some to smaller clusters.

#### I.2 Outliers After ERRASER Correction?

In order to asses the extent to which ERRASER corrected suites remain outliers, we apply our MINTAGE Algorithm 2.1 to the ERRASER training data set 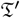 using the tuning parameters from Supplement G.2.

As shown in Figure 26, this returns 15 clusters. The largest corresponds to the A helix shape as in Figure 20, but is even more dominant here. The overall clustering is comparable to that from Figure 20, in particular the four clusters displayed in the respective right panels are similar (Clusters 1, 2, 5, 7 from the benchmark data set in Figure 20 roughly correspond to Clusters 1, 2, 3, 8 from the ERRASER training data set 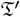 in Figure 26). Table 2, however, shows that approx. 80 % of the clash suites from 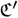 corrected by ERRASER remain outliers.

**Fig. 26:**
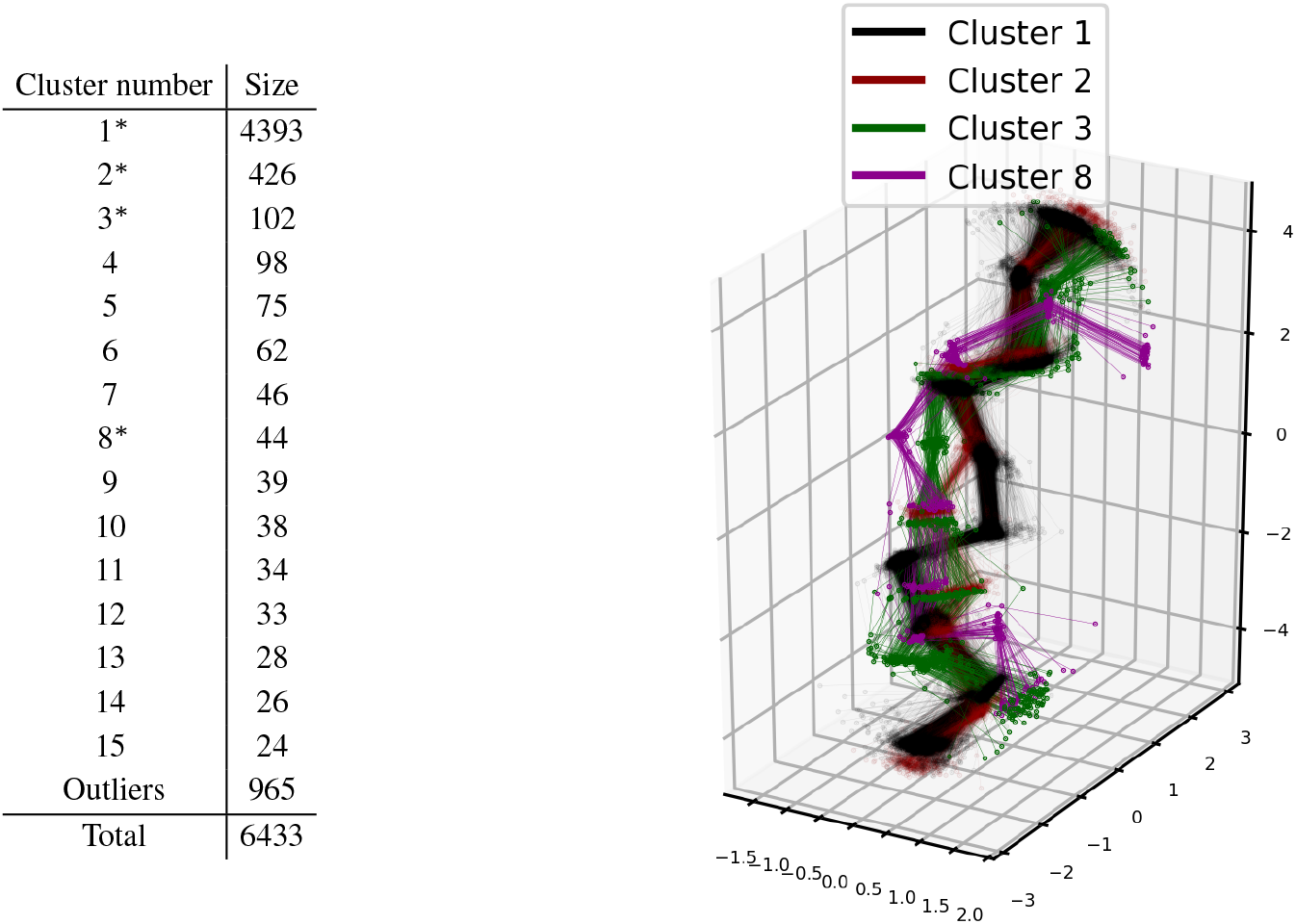
Left: The MINTAGE clusters and outliers (numbers and sizes) of the ERRASER training data set 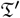. Asterisks mark clusters displayed in the right panel. Right: Five exemplary clusters that can be well displayed together. Notably, Clusters 1, 2, 3 and 8 correspond to the MINTAGE Clusters 1, 2, 5 and 7 of the benchmark data set in Figure 20.

**Fig. 27:**
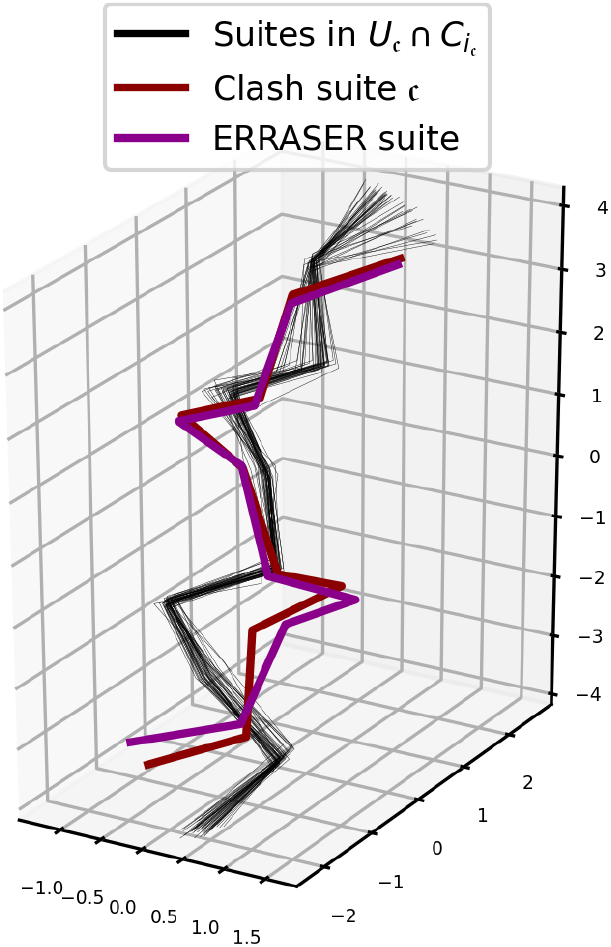
The clash suite 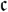 (red) from Figure 8 in the main text and the dominating cluster (black) from the ERRASER training data set 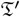 within the 50 nearest mesoscopic shapes in the neighborhood 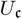 (defined in (1) in the main text), giving their CLEAN MINTAGE correction (in the “middle” of 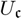, see Figure 8) and the corrected suite proposed by ERRASER (magenta).

**Table 2.** Numbers of clash suites corrected by ERRASER into MINTAGE clusters (numbered as in Figure 26) of the ERRASER training data set 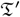, out of the 73 clash suites forming the ERRASER test data set 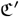.

| Cluster number | number of ERRASER corrected clash suites |
| --- | --- |
| 1 | 4 |
| 2 | 4 |
| 5 | 1 |
| 6 | 4 |
| 11 | 1 |
| 15 | 3 |
| Outliers | 56 |
| Total | 73 |

The exemplary Figures 10 in the main text and 27 here illustrate the reason: ERRASER performs clash correction locally and moderately and in consequence, its results often seem too restrictive. Guiding correction by the mesoscopic scale, as in our CLEAN MINTAGE backbone correction method, results in, often only subtle, changes on larger scales, which are often less restrictive on the suite scale. Indeed, this leads to correcting clash suites into previously classified shapes, i.e. similar to clash free suites in the training set.

## Footnotes

1 The validation tools in PHENIX, see Liebschner *et al*. (2019), reports a clash if van der Waals shells overlap by more than 0.4 Å, see Word *et al*. (1999).

